# Ecological impacts of poultry waste on urban raptors: conflicts, diseases, and climate change implications amidst pandemic threats

**DOI:** 10.1101/2023.07.13.546415

**Authors:** Nishant Kumar

**Affiliations:** Dr. B R Ambedkar University of Delhi, Lothian Road. Kashmere Gate; Delhi - 110006; Wildlife Institute of India, Post Box # 18. Chandrabani. Dehradun - 248001; Edward Grey Institute of Field Ornithology, Department of Biology. 11a Mansfield Road. University of Oxford.OX1 3SZ.; Mansfield College, Mansfield Road. University of Oxford. OX1 3TF.

**Keywords:** Food security, Poultry production and distribution network, Ethno-ornithology, Waste ecology, Scavenging, Ecosystem Service, Zoonoses

## Abstract

The dramatic increase in poultry production and consumption (PPC) over the past decades has raised questions about its impacts on biodiversity, particularly in the Global South. This study focuses on the ecological and environmental impacts of PPC waste metabolism at Asia’s largest livestock wet market, located next to the continent’s largest landfill of *Ghazipur* in Delhi, which I have been monitoring since 2012.
Daily processing of >100,000 poultry-fowls at *Ghazipur* results in an annual production of ∼27,375 metric tonnes of poultry-waste, attracting massive flocks of Black-eared kites, migratory facultative scavengers that winter in South Asia. Approximately >33,600 kites foraged in the area every day and disposed 8.83% of the total PPC slaughter-remains produced during October-April. However, with their return migration to Central Asia, kite flocks over *Ghazipur* reduced by 90%, leading to a proportional decrease in scavenging services. Absence of kites from the larger, migratory race during May-September did not elicit any compensatory response from the small Indian kite, whose numbers over landfill remained unchanged. This raises vital questions about microclimate impacts by green house gases (GHG) released from massive amounts of routine detritus. Bearing in mind the prevalence of ritual feeding of meat chunks to kites in Delhi, my research indicates how life-history traits (migratory vs. resident) enable exploitation of specific anthropogenic resources, creating distinct kite-niche(s). Other opportunistic scavengers, e.g., dogs, rats, cattle-egrets, several passerines, and livestock (fishes and pigs) also benefited from PPC waste.
Public health and ethical concerns, including Avian-influenza outbreaks in 2018-21 and pandemic-lockdowns from 2020-22 - that affected informal meat processing - reduced the flocking of kites at *Ghazipur* by altering spatial dispersion of PPC remains.
Waste-biomass driven cross-species associations can exacerbate zoonotic threats by putting humans and animals in close contact. The ecological impacts of waste-based biomass, as well as the aerospace conflicts caused by avian scavengers that cause birdstrikes must factor in the integrated management of city waste. The quantity, type, dispersion, and accessibility of food-waste for opportunistic urban fauna in tropical cities along avian migratory pathways are crucial for public health, and for conservation of (facultative) migratory avian-scavengers like Eurasian Griffons and Steppe Eagles that are facing extinction threats.

**Lay Summary:**

- The global trend of increasing consumption of broiler chickens, driven by rising incomes in tropical cities, has significant ecological implications for both native and migratory birds, as well as other commensal species.
- The resulting large amounts of debris produced by poultry production and consumption have created a “chicken reconfigured biosphere” in cities along migratory paths.
- To better understand the local and global impacts of poultry production and consumption chains, I conducted a long-term study at Asia’s largest livestock wet market in *Ghazipur*, Delhi.
- The findings reveal that informal handling of poultry waste and cultural practices have had significant impacts on animals that scavenge on the slaughter remains, particularly during the bird flu and COVID-19 pandemics.
- The study recommends ways to minimise conflicts and health risks and reduce the potential impacts of rotting garbage on the climate by accommodating animals that have adapted to shared urban environments.

## Introduction

Over the last six decades, the production and consumption of poultry have dramatically increased, leading to the current situation where the overall biomass of poultry-fowl now exceeds the combined value of all wild birds on Earth by more than 300% (Bar-On et al., 2018; Bennett et al., 2018). Despite the overwhelming evidence that modern global food systems are amongst the primary drivers of defaunation (Bennett et al., 2018; Tscharntke et al., 2012), the ecological impacts of poultry intensification on biodiversity remain largely overlooked, particularly in regions such as South Asia, Africa, and Southern China, where urbanisation is rapidly increasing, and per capita demand for poultry meat by 2030 is projected to rise by approximately 200% (FAO, 2022). While urbanisation has been associated with growth and innovation, and scaling in cities (Bettencourt et al., 2007), it is critical to monitor the massive ecological and environmental impacts of a global standing crop of >25 billion broiler-fowls beyond laboratory-centric studies that primarily focus on carbon footprint and antimicrobial resistance (Bennett et al., 2018). Ecologists are yet to fully examine the far-reaching consequences of poultry intensification on biodiversity, a knowledge gap that must be addressed in the interest of global food security, environmental health, and biosecurity (Gržinić et al., 2023; Tscharntke et al., 2012).

Public health concerns related to poultry production and consumption (PPC) often centre on bacterial or viral contamination of broiler birds (e.g., Hassell et al., 2019; Khan et al., 2020; Woolhouse et al., 2015). In developed regions, there is widespread adherence to scientifically informed poultry management practices that are incorporated from farm-to-pot, to address disease concerns emanating from industrialised PPC (see Thieme, 2013). However, ecological and, thereby, health implications of poultry intensification in developing regions remain poorly understood (Pattison, 2008; Vaarst et al., 2015). This knowledge gap is particularly concerning given the existing skew in the field of urban ecology, with most studies being based in western cities (e.g., review in Marzluff, 2017). Livestock-driven biosphere reconfiguration by humans is a significant challenge due to the ecological and environmental impacts of animal food production, as noted by Dopelt et al. (2019). To put the ecological impacts of more than 65% loss and wastage of food in the meat sector at the consumption stage in perspective (Karwowska et al., 2021), this study focused on the metabolism of food waste from the poultry sector in a developing world setting.

Studying the ecological and environmental consequences of meat-detritus is critical for several reasons. ***First***, developmental heterogeneity in many countries within the Global South affects the possibility to maintain uniform and widespread social rationality and planning for solid waste disposal. Therein, people frequently rely on informal channels to rid of waste, including non-human agencies (Statistica, 2023). ***Second***, such developing countries are experiencing maximum urban growth, and most of the upcoming megacities will be in these regions (Bourdeau-Lepage & Huriot, 2007; UNO, 2018). Consequently, human refuse within urbanising tropics has seen exponential growth, e.g., India’s national capital, Delhi, has the highest per capita production of waste in the country. The national capital has overseen a 300% increase in the amount of solid waste over the past 20 years (S. Kumar et al., 2017). The organic constituents of such enormous urban refuse that afford numerous commensals with vital food-subsidies are predictably dispersed within heterogeneously developed, poorly planned tropical cities. Such opportunistic food resources, whose access to urban animals are patterned in space and time (e.g., Griffin et al., 2017; Oro et al., 2013), are a significant ecological concern within a contested and finite city space (see Griffin et al., 2022; Kumar, Jhala, et al., 2019; Wheat et al., 2019; details in the methods section). ***Third***, a recent study by the National Family Health Survey, India (Paswan et al., 2016), found the growth in per capita income to be closely linked to progressive enhancement of per capita meat consumption by citizens (Roser, 2013; Satterthwaite et al., 2010). Among various Indian states, Delhi, reportedly, has registered the maximum increase in its per capita consumption of meat over the last decade. This rise in meat consumption can be attributed to the increase in the city’s per capita income, which is more than thrice the national average (Devi et al., 2014; Paswan et al., 2016). ***Finally***, a combination of above factors, including concentrated poultry consumption in urban areas, creates a geography of opportunistic responses to the enormous slaughter-biomass by facultative scavengers like kites, crows, dogs, and rats often associated with meat-shops, garbage and abattoirs (Kumar et al., 2019). Despite the significance of how multiple human and non-human stakeholders interact with poultry, from production to consumption, and the overall culmination of food waste, only a few authors have incorporated human socio-economic factors as an integral component of their research (see Faraji Mahyari et al., 2021; Gerber et al., 2008; Guèye, 2000; Mozhiarasi & Natarajan, 2022).

In recent years, numerous studies have focused on the impacts of human infrastructure, such as overhead transmission wires, wind energy farms, and artificial lights at night (ALAN), on migratory avifauna (Drewitt & Langston, 2008; Drewitt et al., 2021; Horton et al., 2019; Nilsson et al., 2021). However, few have explored how the availability of predictable foraging resources at garbage dumps, landfills or abattoirs affects the movement and behaviour of migratory bird populations (e.g., Gilbert et al., 2016; Kumar et al., 2020). This is particularly concerning given the massive growth of meat waste in tropical areas, and when cities/towns producing this waste are frequently located along migratory paths (Galbraith, Jones, Kirby & Mundkur, 2014). Additionally, we have a limited understanding about how the amount and spatial dispersion of urban refuse affects human-animal social-ecological relationships. Gangoso et al., (2013) described one such relationship between the people of Socotra, Yemen, and Egyptian vultures *Neophron percnopterus*, but we need more research to fully understand these complex interactions (see Kumar et al., 2019). Ultimately, the proliferation of poultry and other urban animals can significantly alter ecosystems and sanitary services, affecting human-animal coexistence. Despite such potential cascading eco-evolutionary effects, little research attention has been given to these issues (Oro et al., 2013).

Urban ecology - a discipline encompassing aforementioned issues - is a rapidly growing domain, but researchers have largely ignored the ecological impacts of waste-based biomass. While some recent studies have examined the effects of waste on wild birds and mammals frequenting garbage points and landfills, there is a notable lack of research on the topic (Gil-Fernández et al., 2020; Katlam et al., 2018; Plaza & Lambertucci, 2017; Prange et al., 2003; Vuorisalo et al., 2014). This may be due to the relatively organised solid waste management practices in many Western countries, where urban ecology research has been most prominent (Marzluff, 2017). Additionally, there is a lack of mechanistic explanations as to how access to waste-based food subsidies mediates ecology and behaviour of urban animals (see Kumar, Gupta, et al., 2018; Kumar, Qureshi, et al., 2018). Thus, the findings of my longitudinal study since 2012 on the ecology and metabolism of poultry waste in Delhi’s *Ghazipur* - Asia’s largest livestock wet market situated adjacent to the city’s largest landfill - shed light on complex interplays between multiple organisms in urban environments, addressing important gaps. The study holds special relevance in the wake of recent political discourse surrounding the management of landfills in Delhi. Various political parties in their representations included the flattening of these waste units as part of their electoral mandates and agendas (BJP Manifesto, 2022; K. Sharma, 2022). However, it is crucial to recognise a significant oversight in treating these landfills merely as inert hills composed of garbage and construction waste. The organic matter present in these landfills, particularly the unsegregated waste containing meat detritus, plays a crucial role in supporting and sustaining a distinct community of opportunistic commensals, such as kites, dogs, rats, and even livestock that urban poor rely upon for their sustenance (see Hendrix et al., 1986).

In this study, I explored complex multi-organismic interactions that are driven by organic detritus in urban environments. By examining the chronology of seasonal, and ‘gradual vs. stochastic’ changes to the availability of meat waste, such as those caused by Avian influenza and COVID-19 pandemic lockdowns, I could afford quasi-experimental insights into how the biotic and abiotic components of landfill(s) represent a living ecosystem (Plaza & Lambertucci, 2017), wherein, fluctuations in food-subsidies may impact the movement and habitat choices of both resident and migratory fauna, e.g., kites in Delhi. The manuscript argues that a poor understanding of the ecological underpinnings of urban organic waste hinders progress in several areas, including the incorporation of ethical practices in poultry transport and processing at *Ghazipur*, municipal solid-waste management to prevent the release of GHG/the proposed flattening of landfills, zoonosis prophylaxis for public health issues, and wildlife management to ensure safe aerospace for both birds and aircraft.

## Materials and methods

### Study Area and the ecology of poultry waste

Delhi is a megacity of more than 29 million inhabitants, currently covering an area of 1500 km^2^, which is overlooking constant, rapid infrastructural expansion (*Census 2011 India*, 2011; UNO, 2018). It is polycentric and heterogeneous, with a multitude of urban configurations, which make it difficult to establish a linear urban-rural gradient. The climate is semi-arid, with a mean annual precipitation of 640 mm, mainly concentrated in July and August during the monsoon season. Temperature ranges from a minimum mean value of 8.2°C in the winter to a maximum mean value of 39.6°C during the summer (IMD, 2022). The vegetation of the general region falls within the ‘northern tropical thorn forest’ category (Champion & Seth, 1968).

The poultry production and consumption (PPC) network, which encompasses poultry production, transport, and processing, before consumption, is intricately associated with select set(s) of people from a variety of socio-economic backgrounds, like farmers, butchers, transporters, traders and cleaners, and animals. I focused on PPC links within waste processing subsystems, that is composed of humans as well as non-human agencies. Delhi, as a representative city for South Asian urbanism (Shahmoradi, 2013), has three characteristic aspects that significantly shape PPC in the region. ***First***, like other solid waste disposal systems in the city, the informal handling and segregation of PPC waste generates informal livelihoods (see Kumar et al., 2019 for details). Garbage piles of varying sizes and contents, linked with informal labour as the primary agency for processing waste, generates spatially patterned and close interactions between urban poor and opportunistic commensals such as stray dogs, rats, cattle, and various facultative avian scavengers (kites, crows, egrets, and pigeons). Owing to their similarities in urban heterogeneity, tropical cities in the global south have widespread congruence over treatment of urban detritus. This detritus and associated animals in shared living spaces give such tropical cities, a representative visual and olfactory character. With urban poor at their core, such communities on city detritus collectively represent the stench and sensibilities of tropical urbanisation (Doron, 2021).

***Second***, PPC in Delhi and nearby regions is facing significant changes due to upheavals in human meat consumption that generate subsystems of animal waste. These changes are primarily driven by the collective impacts of demand-supply chains, urban planning, and poultry management on public health concerns (GoI, 2022; S. Kumar et al., 2017). As a result, the traditional practice of rearing poultry in sheds and backyards is being replaced by organised poultry cooperatives (GoI, 2019), which is affecting the number of country-breeds and homogenising the overall poultry germplasm (Bennett et al., 2018; Chatterjee & Rajkumar, 2015; M. Singh et al., 2022).

***Third***: aforementioned changes, however, are constrained by the increasing volumes of unsegregated waste, animal remains and excreta, and the lack of suitable new places for garbage disposal, leading to conflicts of management among various civic bodies. The Delhi Development Authority is responsible for making the city’s Master Plan every 20 years, including spaces for waste, while municipal corporations are responsible for garbage collection and dumping spaces (Bettencourt et al., 2007; Talyan et al., 2008). The Department of forest and wildlife, under the Government of National Capital Territory of Delhi is addressing encroachment by unauthorised dumpers of waste, and animal welfare organisations advocate for improved management. However, the failure to find new sites has resulted in all three of Delhi’s landfills continuing to function more than a decade after court orders for their decommissioning were passed. Additionally, several sub judice ‘Public Interest Litigations’ filed by animal welfare activists have enforced a complete ban on formal processing of poultry at *Ghazipur*, the largest market facility in Asia and the location of the continent’s largest landfill. The location also receives offal from various livestock meat processing units in the vicinity. This ban has led to the decommissioning of poultry slaughter on the grounds of animal cruelty, which stochastically altered the spatial dispersion of PPC waste as food-subsidies to urban fauna since September 2018. Given the likelihood of further disruptions to where and how poultry can be reared and consumed, new regulations are already in place with respect to how landfill sites can be used for the disposal of slaughter remains (details in Kumar et al., 2019).

The ongoing political discussions regarding the flattening of landfills in Delhi have sparked competing agendas among different political parties, advocating for the repurposing of waste materials for road construction and electricity generation (BJP Manifesto, 2022; K. Sharma, 2022). This shift in focus on waste management has emerged as a significant aspect of election manifestos in the context of a tropical megacity. However, upon careful analysis of all available manifestos, a striking observation is the apparent absence of consideration for potential ecological implications and challenges associated with the rapid dismantling of landfill (BJP Manifesto, 2022; K. Sharma, 2022). While the concept of flattening mega landfills and utilising waste materials for construction and energy production holds promise in terms of reducing environmental burdens and generating sustainable resources, it is imperative to approach these initiatives with caution. Therefore, for this manuscript, I attempted a comprehensive assessment of feasibility and potential risks from PPC and landfill detritus to avoid unintended ecological consequences. Specifically, I gave careful attention to the organic composition of the waste, encompassing various organic material and meat detritus from PPC/cattle/fish processing units, in order to mitigate potential direct and indirect impacts.

### Human and urban parameters of poultry waste

This study is part of a comprehensive, ongoing research project on the ecology and ethno-zoology of urban scavengers in Delhi, which began in 2012 (Kumar et al., 2019). Through regular surveys at 32 sampling plots spanning approximately 1 km^2^ in a stratified-random design, my field sampling has systematically covered a range of urban settings, from semi-natural to extremely built-up sites, including all three sanitary-landfills (for more information, see Kumar et al., 2019). Given that urbanisation is an ongoing process in Delhi, which has created a melting pot of cultures with people from across the Indian subcontinent (Bhagat & Mohanty, 2009), this study examined the ecology of non-human and urban parameters of poultry waste. In addition to monitoring the metabolism of poultry waste and ecology of opportunistic scavengers (Pickett et al., 2016), I employed a Delphi-like ethnographic approach to understand the range of stakeholders and their interactions with poultry waste and facultative scavengers (Esmail et al., 2020; Mukherjee et al., 2015).

During my field visits to Delhi’s landfills, I conducted ethnographic research that involved interactions with an average of 27.03 new onlookers (SE: 2.31) within each of the three sampling units. These individuals willingly engaged in conversations about the impact of poultry detritus, and I employed various methods, including iterated surveys, facilitated discussions, structured elicitation, and aggregation of individual perceptions, to gather direct input on the importance of poultry-based food subsidies for opportunistic scavengers in the city. To address individual-level psychological biases, I interacted with new, randomly selected respondents from the same stakeholder unit, and thematically organised their responses to identify correlations in socio-cultural and professional backgrounds (Esmail et al., 2020). Additionally, I collected information as to how the prevalence of poultry detritus links with opportunistic scavengers, such as dogs, rats, other avian commensals, and livestock, under the purview that meat detritus could mediate prosocial or agonistic human and non-human animal interactions in shared spaces (see Moleón et al., 2014). These observations allowed me to understand how informal and state-driven human agencies formed complex but predictable spatio-temporal interactions with native and invasive animal agencies flourishing on waste (Kumar et al., 2019).

### Urban scavenger’s guild on landfill detritus: estimating the number of kites on the landfill

Black Kites are frequently seen flocking over landfill sites in South Asia, where they congregate in numbers ranging from a few hundred to several thousands, scavenging on detritus from kitchens, eateries, and slaughterhouses (Kumar et al., 2020; Fig. 1). At landfills like *Ghazipur*, it is almost impossible to visually estimate the number of kites in the region using conventional methods like point points, or through total counts from a single point location. Apart from hovering over the dump these huge kite flock also forage at multiple locations by perching on the garbage. Therefore, it is essential to count the number of birds in flight and those perched on the garbage to arrive at a realistic estimate. To estimate the number of these birds hovering over, and perched on the landfill, I conducted a systematic survey from ten vantage points with good visibility. These vantage points were selected using a systematic sampling method in 2012-13 that I describe below.

**Fig. 1.**
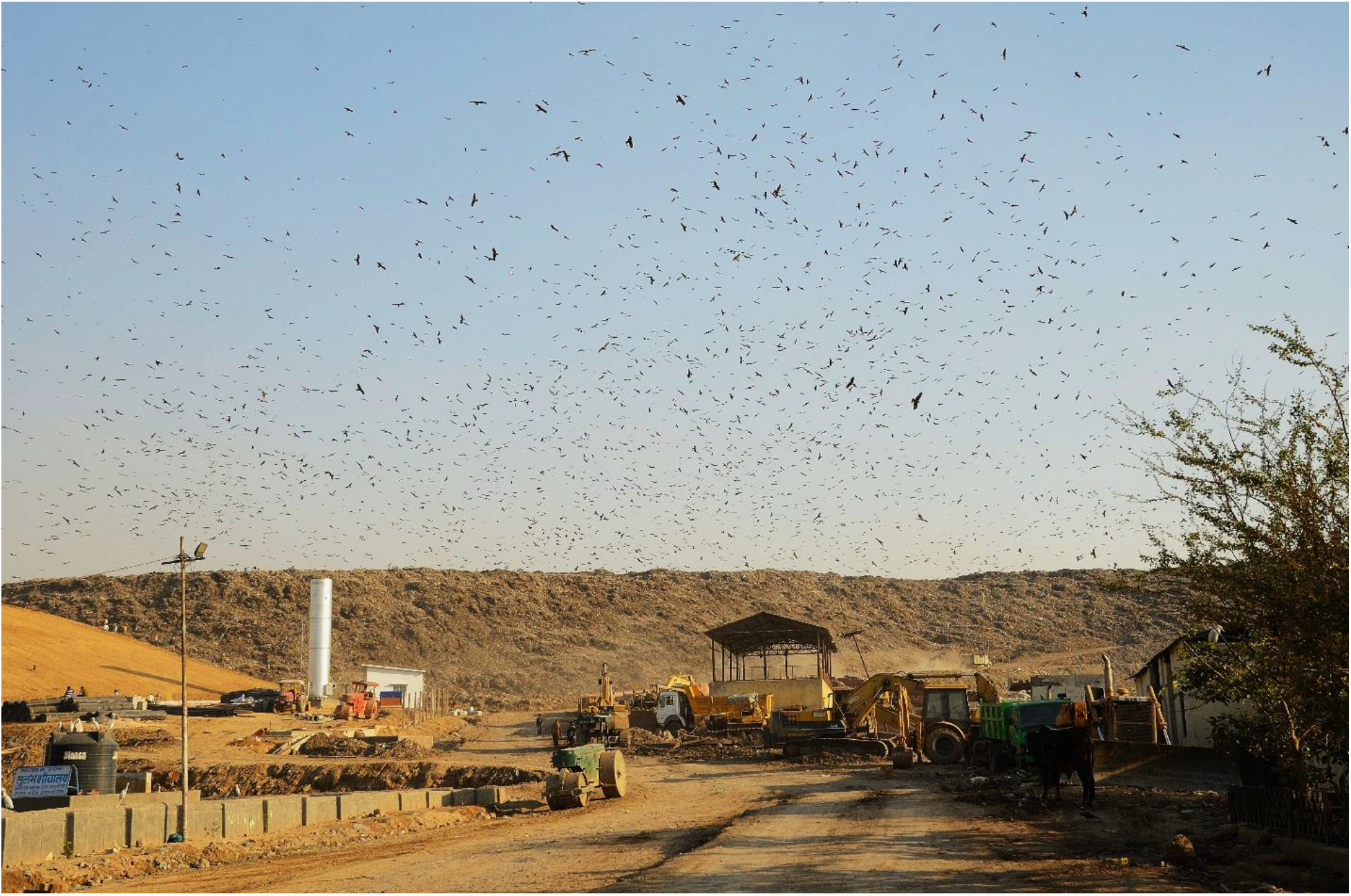
Massive flocking of Black-eared kites *Milvus migrans lineatus* is typical for landfills within tropical cities across South Asia. This image from February 2013 depicts a beholding view of >10,000 kites that regularly congregate over and around the *Ghazipur* landfill spread over 70 acres. Kites and other commensals mostly capitalise over poultry processing waste at and from *Shaheed Ashfaqullah Khan* Chicken, Fish and Egg market (SAK) (Government of National Capital Territory of Delhi) in the vicinity. (Photo Credit: Fabrizio Sergio).

The number of kites in the region varies significantly throughout the day, depending on the behaviour of these soaring raptors, which utilise air thermals for flight and foraging. The human detritus found in the dump, primarily from poultry/livestock slaughter, attracts a wide range of avian and other scavengers, including opportunistic feeding of livestock by the urban poor from nearby informal establishments, like dairy farmers (Kumar et al., 2019; see Fig. 2 for a visual representation of on-ground situations and the strength of kite flocks). To determine the time of maximum congregations over a 24-hour diel cycle, I conducted repeated counts and photo shoots of the dump area from a fixed vantage point every hour between 0500 hours and 1830 hours (the end of daylight) from 2012-14. I used two steps to estimate the number of kites - total counts and the estimation method using an open access Java based ImageJ software (https://rsbweb.nih.gov/ij/), as described in detail by Kumar (2013). By obtaining the mean of the numerical estimates from the total counts at the vantage points, I arrived at an estimate for the number of birds in flight.

**Fig. 2.**
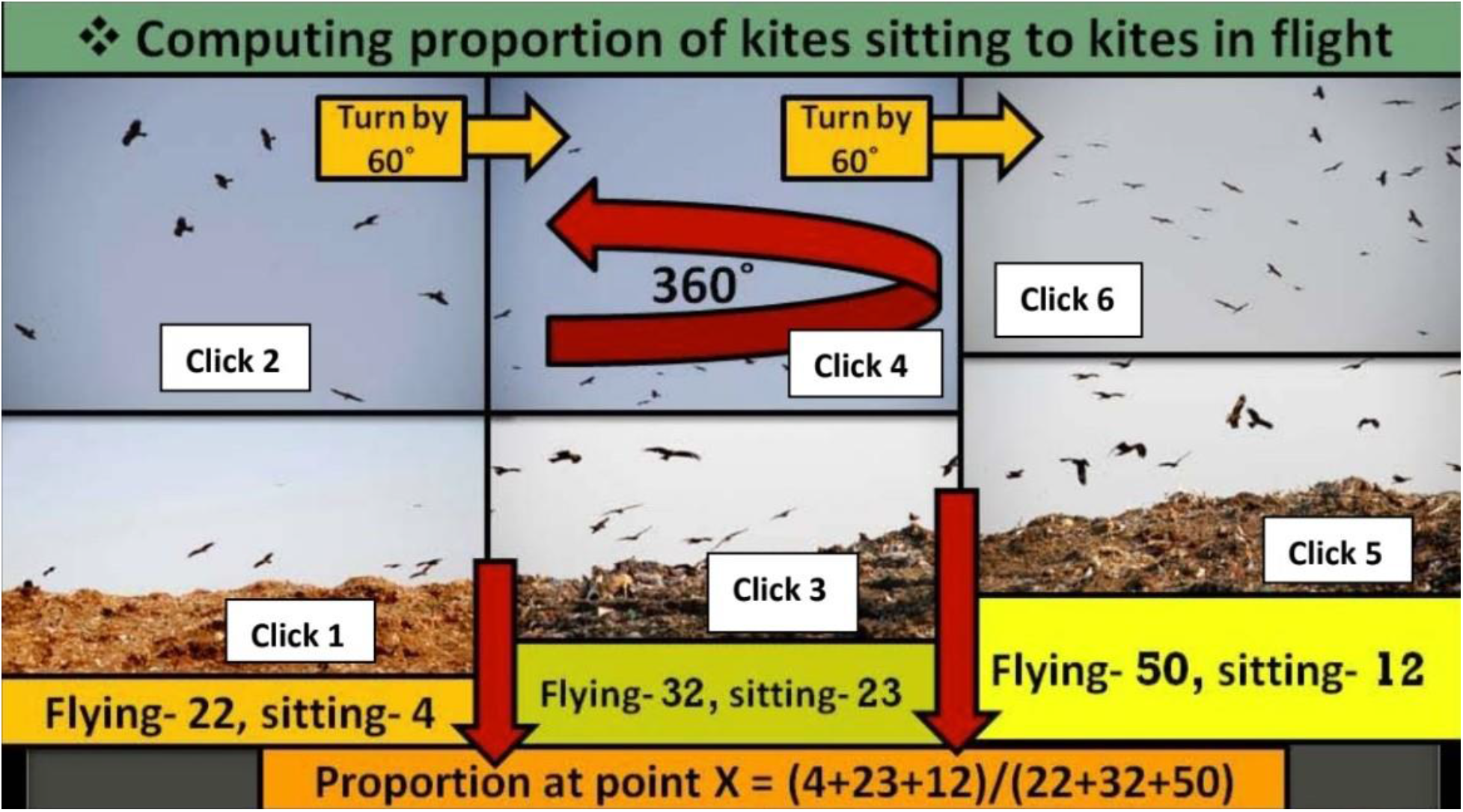
To count the black kites at *Ghazipur*, I stationed 10 people, each with a camera, at predetermined locations in the area during 2012-14 (Figures 1 and 2). Each person first took a picture of the kites sitting on the dump, then tilted the camera upwards to take a picture of the kites in flight (Figure 3). They then turned 60 degrees and repeated the steps, being careful not to overlap any of the previous frames. The person at point ‘X’ continued this process, turning 60 degrees each time, until they had captured a 360-degree view of the kites in five passes. Finally, they took an overhead shot of the kites flying overhead.

Subsequently, to obtain the mean value for the ratio of **“*kites perched on the dump: the kites in flight*”**, I used Image J and manual counts to estimate kite numbers from the photographs that were clicked under a schematic, as described: I stationed 10 people, each with a camera, at previously chosen systematic locations in the regions during 2012-14 (Fig. 1 and 2). Each person first clicked the kites sitting on the dump followed by tilting the camera upwards to shoot the kites in flight (Figure 3). This was followed by a 60° turn and repeat of the earlier steps, while attempting to avoid overlaps with the previous frame. The concerned person at point ‘X’ continued this with successive 60° turns till s/he completed capturing kites in his/her 360° view through 5 replicates of the initial step. In the end, kites flying over his/her head were captured with an overhead shot. Thus, on an average, 6 clicks of kites perched in the region and the 7^th^ click of the kites that were in flight afforded us the ratio of completed shooting of kites available for a person from a given point on dump. Rest nine persons also simultaneously followed the same steps at their respective points. This activity was synchronised using cellular phones to avoid double counts of birds as they change locations in a span of a few minutes. To estimate the minimum number of kites benefiting from the poultry detritus in a 24-hour period, I used telemetry data from GPS-tagged Black-eared kites (14 adults and five pre-adults) that were monitored between 2014 and 2018 (details in Kumar et al., 2020).

**Fig. 3.**
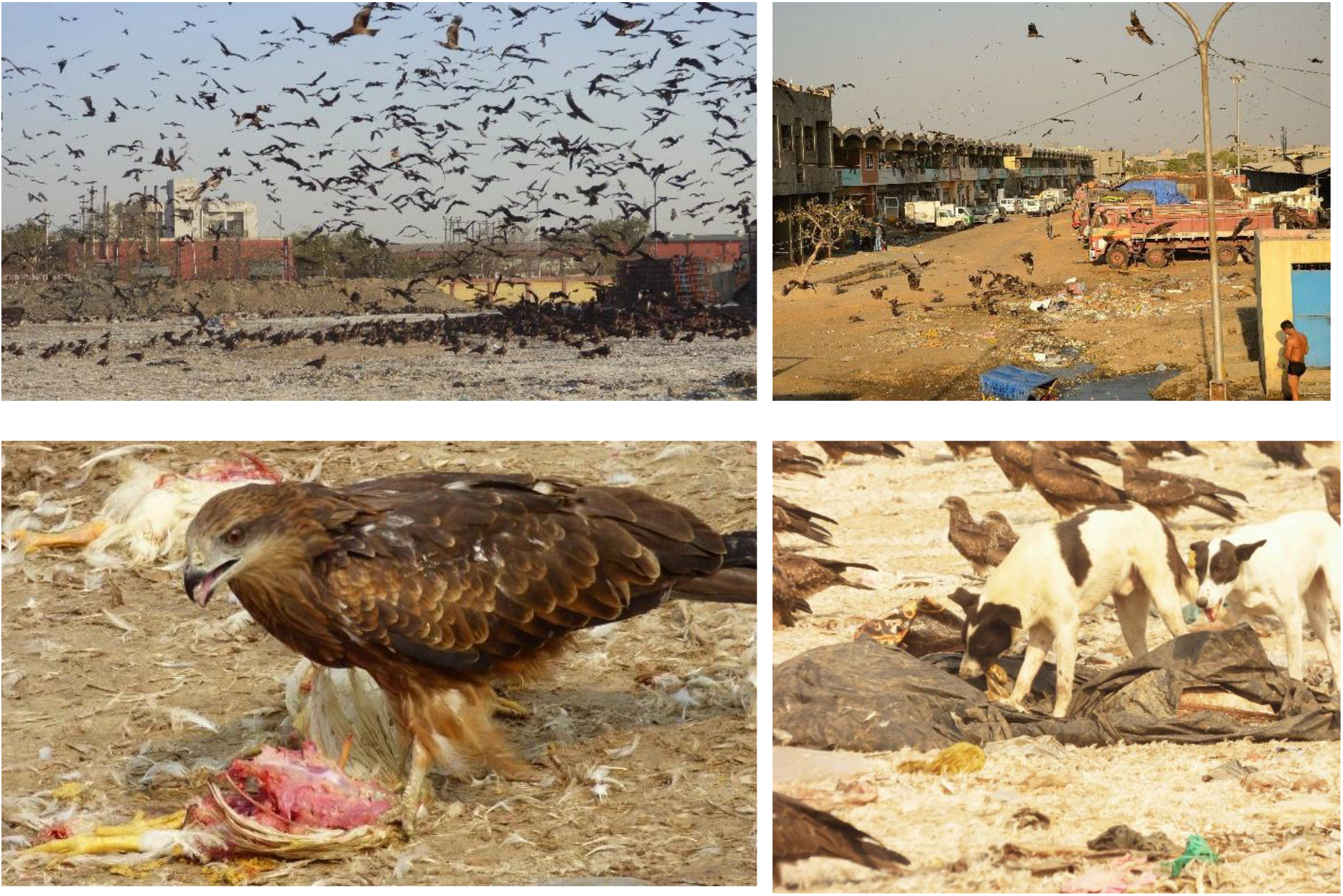
Food scraps from poultry slaughter provide opportunities for interactions between humans and animals that scavenge on garbage piles. At landfill systems like *Ghazipur* that are associated with meat processing units, this creates systematic relationships between different species, which are poorly understood. The forms and functions of these guilds, which develop over urban food and foraging resources, affect how people view animals that live in cities and how animals adapt to the heterogeneous social environment of cities.

### Calculating GHG emissions from landfilling of PPC waste

Based on the estimate of waste that was not consumed by kites, which I assumed the SAK sent to the *Ghazipur* landfill (Singh, 2021), I used the Intergovernmental Panel on Climate Change (IPCC) methodology for estimating CH_4_ emissions from landfilling. These estimates are based on the first order decay (FOD) approach, which was applied with the default parameters and regional specific landfill data in the IPCC waste model, following Ghosh et al. (2019); Yu & Zhang, (2016). The paper shall present quantitative assessments for kite numbers at *Ghazipur* and qualitative assessments for the ethnographic surveys, involving multiple stakeholders in the region. All means are given ± 1 SE.

## Results

Interviews with residents and the workers at *Ghazipur* revealed that the distribution of kites on the dump, and poultry processing and disposal facilities across Delhi underwent significant variations. Since 1992, with the establishment of the National Capital Territory of Delhi, the majority of these facilities have slowly been consolidated at *Ghazipur*, established near the landfill site by relocating livestock processing units from the densely populated areas of the city, including *Idgah* and *Jama Masjid*. The landfill at *Ghazipur* was created as part of preparations for the 1982 Asiad Games to accommodate the waste away from the main city. Similar international sporting events, such as the 2010 Commonwealth Games, have played a pivotal role in shaping Delhi’s infrastructure development and spatial dispersion of waste in the city. In addition to these events, Delhi’s infrastructural growth was also shaped by the central government’s five-year plans and the Delhi Development Authority’s 20-year master plans, which resulted in significant changes to the concept of waste and hygiene in the city, accompanying large-scale gentrification initiatives across the turn of the century.

### Estimates for kite numbers within Ghazipur landfill area and their scavenging services

Kite numbers in *Ghazipur* landfill area (1 km^2^) underwent large scale variations, based on how these soaring raptors take advantage of air thermals for foraging and flight displays. My observations during 2012-14 to understand patterns in arrival/departure of flocks revealed that these flocks typically start building in numbers around 0800 hrs, with the first peak occurring at around 1000 hrs, when air thermals take effect. The second and third peaks occur around 1400 hrs and 1630 hrs, respectively (Fig. 2). Before kites build their flocks at *Ghazipur*, early in the morning (0530 - 0630 hrs), thousands of house crows *Corvus splendens*, egret spp., pigeons *Columba livia*, Red-naped ibis *Pseudibis papillosa*, Egyptian vultures, house swifts *Apus nipalensis*, and several small passerines (myna spp., wagtail spp., Rosy starlings *Pastor roseus*) also occupied the landfill, feeding on detritus/insects. The numbers of crows and pigeons dwindled with the increase in kite numbers, 0800 hrs onwards.

The GPS data showed that tagged kites spent an average of 3-4 hours in the landfill area. The time of day that the tagged birds (n=20) visited the landfill varied depending on the individual, age, and distance of the roosting sites from the landfill. In comparison to adult birds, all pre-adult GPS-tagged birds (n = 6) roosted in proximity of *Ghazipur* (*χ*^2^= 7.01, P = 0.008). Since three peaks in kite congregations were observed within 24 hours in the region, it appears that multiple groups of kites from across Delhi visited the landfill(s) and foraged for about 3-4 hours before returning to their roosting locations. These observations were consistent throughout the sampling period, which the research team used to plan trapping of birds for GPS tagging. The team found that the birds congregated most often around 0930 hrs, 1330 hrs, and 1630 hrs. Multiple team members counted kites in flight at the nearby Ghazipur and *Shaheed Ashfaqullah Khan Chicken, Fish, and Egg market* (SAK) during peak congregation times. Based on these repeated visual counts, I estimate that there are approximately 10,000 kites. Based on this number, I estimate that at least 33,600 kites feed on poultry detritus at *Ghazipur* landfill every day. I arrived at this figure by combining the total number of kites in flight with the number of birds perched on the landfill and nearby. The number of kites in flight was estimated by multiplying the number of major flying congregations (10,000) by 3. The number of perched birds was obtained from the overall ratio of ‘birds in flight: birds perched’ within *Ghazipur* area, using the equation from (N. Kumar, 2013). This gave a total of 33,600 kites.

At poultry shops within the SAK facility, procurement, processing, and slaughter detritus management practices were community-specific, associated with social factors such as religion, caste, and class. For instance, at the SAK facility in *Ghazipur*, the majority of workers who handled poultry waste, as well as allied staff and owners/workers at roadside eateries, came from Muzaffarpur, a district in the Indian state of Bihar. During the study period from 2013 to 2022, the SAK facility procured and processed more than 100,000 broiler chickens every year each day, which yielded approximately 400 metric tonnes of poultry meat for local consumption. Poultry fowls were sourced from farms/villages in neighbouring states, such as Haryana, Uttar Pradesh, Punjab, and Rajasthan, and transported to SAK on over 200 trucks/day. The processed meat was sold to local markets, restaurants, and also the aviation industry for use in in-flight meals.

The annual estimate for poultry waste from *Ghazipur*, which includes feathers, skin, heads, blood, and entrails, was approximately 27,375 metric tons. This estimate was based on an average of 250 grams of slaughter remains per kilogram of bird weight (mean weight estimated per fowl = 3 kg). During the winter migration period (October to March), migratory Black-eared kites from central Asia scavenged to dispose of about 8.83% of the slaughter detritus. However, during the April-September period, the birds that remained on or near the landfill (n <1000) could only scavenge and clear less than 0.03% of the total estimated 13,688 metric tons of slaughter detritus. In other words, the Black-eared kites were able to scavenge and clear a significant amount of poultry detritus during the winter migration period, but with their return migration, this ecosystem service decreased significantly during the April-September period. The absence of the larger, migratory kite species during May-August did not cause the smaller Indian kite species, *Milvus migrans govinda*, to increase in number within *Ghazipur* area. The number of birds over the landfill remained unchanged, with an overall bird count of less than 1000 during May-August. Further, the number of kites in flocks at the other two landfills in Delhi was less than 1000, possibly because there are no livestock processing facilities at *Okhla* and *Bhalswa*. People from nearby slums collected poultry slaughter refuse from *SAK* for their own consumption, while farmers used it as feed in nearby pisciculture units within states bordering Delhi. Strong ties between informal and private initiatives helped to ensure that a significant portion of the remaining poultry waste was metabolised through the use of poultry waste in fish farming. Large trucks transported fish from Kolkata (a metropolitan city on the eastern coast of India, 1600 km from Delhi) to SAK, and on their return journey, they loaded poultry waste, which was a major source of food for fish farming in the region, primarily for desi mangur *Clarias batrachus*, an invasive catfish in Asia (Fig. 5). However, these waste transport links between Delhi and Kolkata via circular economic channels only operated during the winter season (November-March), due to the limited lifespan of waste transported by road, which could rot and cause problems. The journey from Delhi to Kolkata took 2-3 days, during which, drivers of transporting trucks faced hostility from local residents along the journey if rotting detritus spread foul smell from the vehicles. The amount of time that waste could be stored before it began to decompose was limited by the amount of time that ice sheets could cover the waste in containers, the reasons it was impossible to move poultry waste to large distances during the summer. During April-October, this waste was either put on landfill or informally collected by regional fish farmers (Fig. 4). In April-May 2018, the government of West Bengal imposed a ban on the entry of poultry detritus from Delhi’s *Ghazipur*. As a result, meat waste from over 100,000 chickens per day was piled up at *Ghazipur* for more than 1.5 months, resulting in more than six-foot-high rotting biomass spread over an area half the size of a football field (Fig. 4). The ban was imposed because of the outbreak of diseases in the fish farms of Kolkata, which was linked with use of Delhi’s detritus as feed. This event highlights the challenges and intricacies of managing food waste and its potential impact on the environment, animals, and public health. The interaction between poultry and fish trade, and their detritus, linked with urban commensals in complex ways. These ecologies of PPC and its detritus, illustrated in Fig. 5, are poorly explored for their implications on human, animal, and environmental health.

**Fig. 4.**
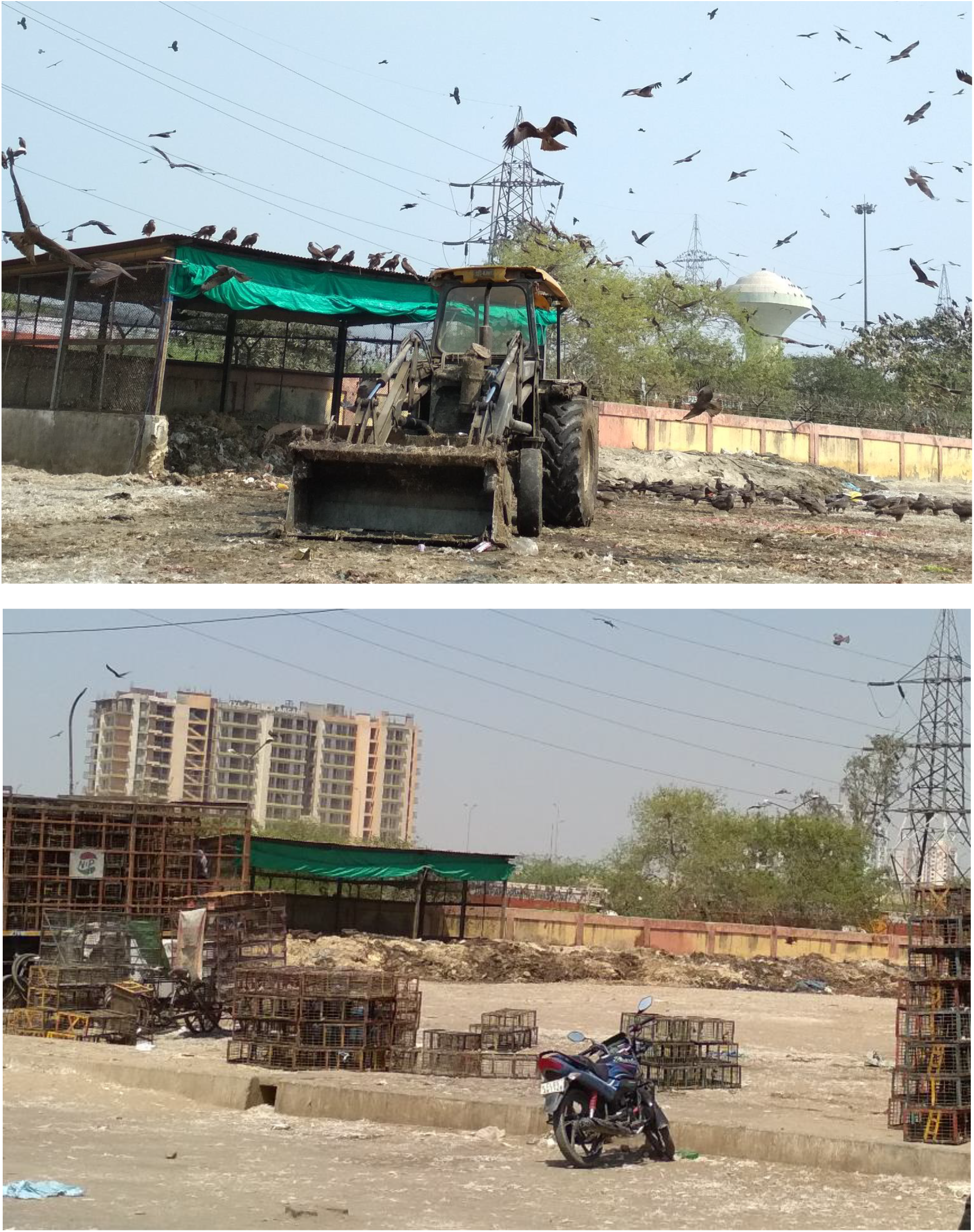
In April 2018, West Bengal’s state government refused to accept poultry waste from *Ghazipur* for its fish farms due to the risk of disease transmission. The waste-to-value chain was disrupted. The pictures show how meat waste from over 100,000 chickens per day was piled up at *Ghazipur* for 1.5 months, resulting in a six-foot-high rotting biomass spread over an area half the size of a football field.

**Fig. 5.**
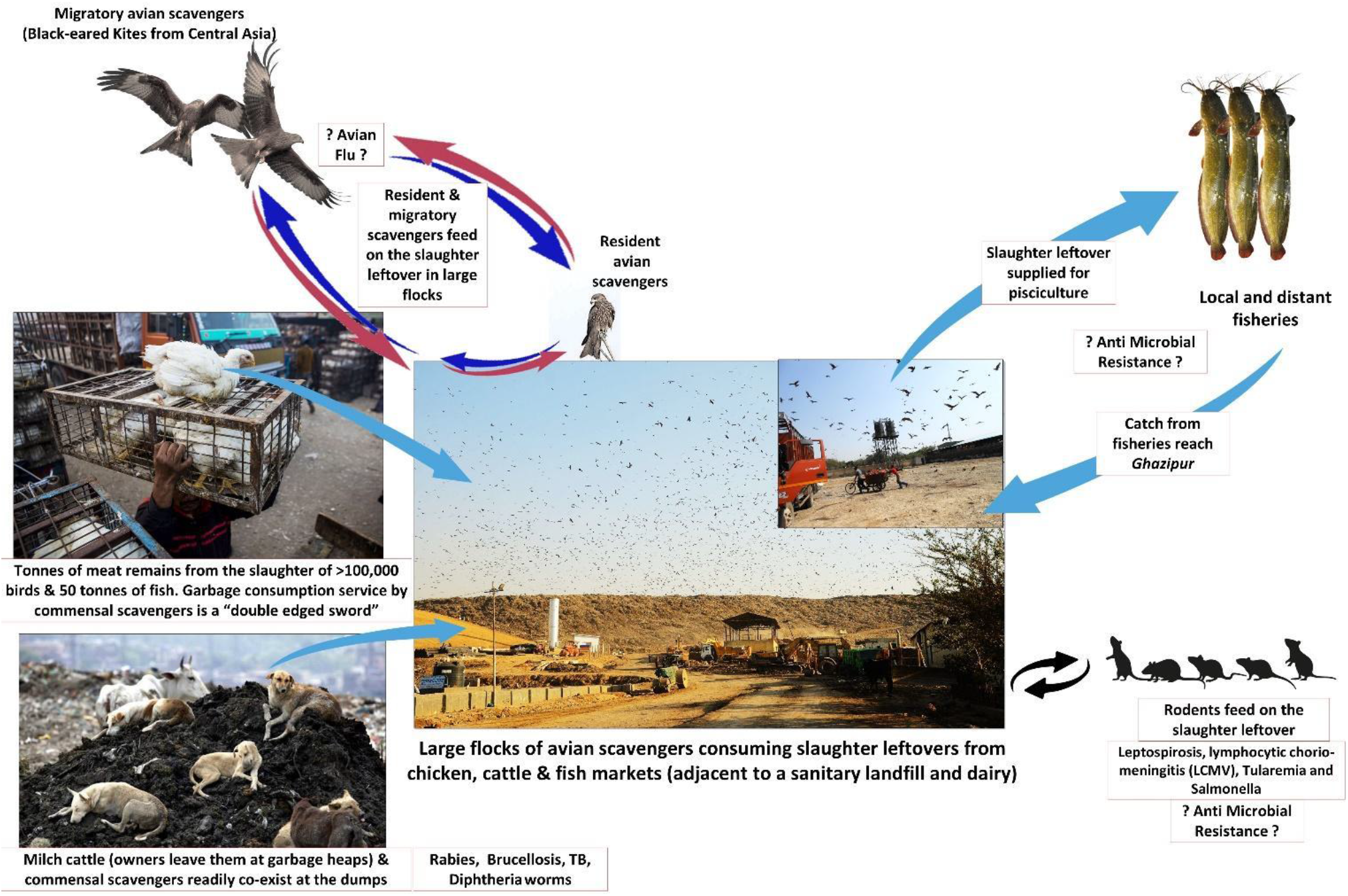
*What goes around, comes around*: the figure shows how people, animals, and waste interact in tropical megacities. It highlights the potential risks of informal handling of meat processing waste, such as bioaccumulation and biomagnification of toxic elements, antimicrobial resistance, and zoonotic diseases. These relationships and their implications are important for understanding the ecological and social dynamics of rapidly changing, human-dominated environments.

### Environmental impacts of meat/organic detritus from Ghazipur landfill: GHG and wastewater

If all PPC waste that is not consumed by kites ends up rotting in landfills (which may not be the case, as dogs, rats, humans, and other animals may scavenge on it), then according to IPCC models, such detritus would produce 17.04 Gg of methane per year. Regardless of how PPC is metabolized (through scavenging, pisciculture, or landfilling), the amount of waste warrants serious consideration from the Municipal Corporation of Delhi. In case we take the landfilled biomass moving from SAK, as estimated by Singh, (2021), then such detritus would produce 2.68 Gg of methane per year.

Furthermore, the wastewater from the poultry slaughterhouse was discharged directly, untreated, into municipal wastewater by formal and informal traders in the region. Such effluents have a high Biochemical Oxygen Demand (BOD) that can harm aquatic fauna. Further, open dumping practices that are prevalent at *Ghazipur* lead to emission of enormous amounts of methane into the atmosphere. The East Delhi Municipal Corporation, in collaboration with the Gas Authority of India Limited and the Indian Institute of Technology, Bombay, launched a pilot program in 2012-13 to harvest methane from the anaerobic decomposition of waste using 40% of the area of the *Ghazipur* landfill. However, the unit was shut down immediately after it began operating. The main reason for this was that the methane harvest was suboptimal due to the mixed nature of the solid waste on top of the landfill.

Poultry detritus presented a significant opportunity for multi-species interactions between native and exotic (migratory) species at *Ghazipur*. During the study, the urban detritus community experienced several changes due to stochastic impacts, such as multiple Avian influenza outbreaks/prophylaxis between 2018 and 2021, ethical and legal issues with PPC in 2018, and the COVID-19 pandemic from 2020 to 2022. In addition, many Hindus refrain from eating meat during the Indian month of *Śrāvaṇa*, which coincides with the monsoon rains in July and August, for cultural reasons. This leads to a regular fluctuation in the availability of PPC detritus. One of the respondents at *Ghazipur* explained:

> “Śrāvaṇa is a time of natural renewal, when the Earth regenerates its vegetation, and the animal kingdom also undergoes reproductive rituals. As a result, Hindus abstain from killing animals for meat, a traditional practice that predates modern animal husbandry and is still practised today.”

Additionally, in Hinduism, it is a common practice to abstain from eating meat on certain days of the week as part of religious observance. Tuesdays, Thursdays, and Saturdays are traditionally considered to be days when meat consumption is prohibited. The availability of poultry waste was affected by monthly and yearly trends, which also had a negative impact on poultry meat sellers, particularly small informal shops, who did poor business during *Śrāvaṇa* or on specific weekdays. Additionally, the Government of the National Capital Territory of Delhi took several preventive measures between 2012 and 2021 by restricting poultry trade in response to avian influenza, further impacting the availability of poultry waste. Improper disposal of carcasses after large-scale culling drives benefited urban scavengers like stray dogs and rats. Tracking kites with GPS tags helped to understand the effects of stochastic changes (such as a ban on chicken slaughter, pandemic lockdown, and avian influenza) on the distribution of food subsidies. However, due to the lack of tagged individuals, it was not possible to assess how food subsidies for rats at *Ghazipur* could be linked to AMR (Fig. 5), despite the existing public health concerns associated with urban rodents.

The decision to ban poultry slaughter at SAK in September 2018 had a significant impact on the point location availability of poultry detritus at *Ghazipur*, which subsequently affected the number of kites congregating on the landfill. Since poultry waste was only available from around 20 informal shops in the SAK area, kites began to congregate at the nearby fish market, as well as on offal found at the landfill and throughout the rest of the city. Counting kites from high points showed that the overall flock size during the peak wintering season was reduced to about one-third of the previous estimate of around 10,000 birds. GPS-tagged kite data also revealed that avian scavengers likely dispersed throughout Delhi after the decentralisation of chicken slaughter in the city, which also caused the dispersion of slaughter waste.

Interviews conducted during prophylactic containment to prevent the spread of bird flu revealed that during the COVID-19 pandemic, a significant amount of misinformation circulated regarding the potential transmission of the disease through poultry consumption, leading to a decline in the consumption of poultry meat and eggs. Thus, poultry was sold at very low prices during the initial pandemic lockdowns from March to September 2020. This misinformation regarding the potential transmission of the disease through poultry consumption spread rapidly through various media channels and social media platforms, often fueled by sensationalised reports and misinterpretation of research findings that examined the survival of the COVID-19 virus on raw meat surfaces. As a result, there was a significant decrease in the consumption of poultry meat and eggs, as people started stigmatising poultry as a potential spreader of the disease. Despite the scientific consensus on the safety of consuming properly cooked poultry products, the misinformation surrounding poultry as a potential spreader of COVID-19 had a tangible impact on consumer behaviour. According to my interviewees, during the pandemic, consumers opted for alternative protein sources from plant origin. This decline not only affected the poultry industry at *Ghazipur* but also disrupted supply chains and posed economic challenges for farmers, informal workers, and businesses involved in poultry production and distribution. The pandemic also led to the creation of satirical songs in regional languages that mocked the situation and spread misinformation surrounding poultry as a potential spreader of the virus. One such song, “*Murga Badnaam Hua Corona Tere Liye*,” means “The rooster has earned a bad name for itself due to COVID-19” (see MCRS Music, 2020).

Market disruption caused the price of chicken and eggs to drop by up to 80% (general drop was at 20%), as people believed that poultry was a potential carrier of the virus. Since major lockdowns in 2020 (April - June) and 2021 (late April -May) coincided with the breeding time of Black-eared kites, when they are in northern Central Asian latitudes, I could not register the impacts of reduced poultry sale on the kite-movements at *Ghazipur*. These changes led to positive shifts in meat consumption by the urban poor who typically cannot afford it. In contrast to Western countries, socio-economic status influenced who consumed the most desirable parts of poultry. For example, the tarsii, skin, head, gizzard, etc., which are typically discarded during broiler processing, were available for Rs. 20/kg (i.e., at 10% of the price of broiler meat) and were typically purchased by people of lower socioeconomic status.

This study provides a clear example of the ecosystem-level impacts of cultivated non-human detritus. This detritus acts as a pollutant and food source, affecting both feral and wild species. Due to human reconfiguration of the biosphere by broiler chickens and other poultry, tropical urban cities like Delhi link society and biota at multiple spatial and temporal scales as an ecological formation. The findings underscore the need to comprehend how animal husbandry activities and practices in developing regions like South Asia connect with socio-cultural, economic, and ecological processes. In most tropical cities, the increasing disposal of solid waste creates enormous foraging opportunities for urban wildlife. However, despite the regional to global impacts of cultivated animal detritus that were discovered in the surveys, its conceptualisation has largely been overlooked by conventional approaches to ecological assessments in cities.

The response of avian scavengers to food subsidies, space, infrastructure heterogeneity, and human agencies showed how these factors interact to create a mosaic of homogenised urban biota. For cities like Delhi, without a rethinking of how the dispersion of poultry detritus is mediated by urban infrastructure and socio-economic factors, investigating its importance as a predictable resource for opportunistic scavengers would have been overlooked. Below, I discuss the implications of these interactions for human, animal, and environmental health. The following are the implications of these interactions for human, animal, and environmental health:

- *Human health*: Avian scavengers can spread diseases to humans through contact with their droppings or by eating contaminated food.
- *Animal health*: Avian scavengers can also spread diseases to animals, such as poultry.
- *Environmental health*: Avian scavengers can contribute to the spread of pollution and the decline of biodiversity.

## Discussion

The article highlights the ecological distinctions between traditional methods of poultry rearing/consumption and modern practices, particularly in the context of rapidly urbanising tropical systems like Delhi. These contrasting aspects underscore the challenges of meeting the growing demand for poultry products in urban areas, while also maintaining traditional production practices that are often associated with rural areas. Furthermore, the article notes that poultry rearing has seen the highest rate of increase amongst all livestock items over the last 50 years (Bennett et al., 2018). However, the majority of poultry reared in South/South-east Asia still happens in relatively rural areas, where backyard farming is a common practice (Gilbert et al., 2015). These findings suggest that there may be an opportunity to promote more sustainable and equitable poultry rearing practices in urban areas, where demand for poultry products is high (Chatterjee & Rajkumar, 2015; Gilbert et al., 2015; Thieme, 2013; Wong et al., 2017).

In addition, field observations pointed out disproportionately higher participation of women in backyard poultry rearing in tropical countries, unlike the distribution and processing networks that are male dominated (noted by FAO, 2019; Njuki et al., 2013; Rota, 2023; Samanta et al., 2018). This gender imbalance along the poultry value chain has important implications for equity, gender-mediated susceptibility to diseases spread by handling poultry, and the economic empowerment of women, and/or poor and oppressed people. Efforts to promote more sustainable and equitable poultry production practices in urban areas must take into account these gender-specific concerns at various stages of PPC value chain (Guèye, 2000; Jensen, n.d.; Njuki et al., 2013; Rota, 2023). My observations provide compelling examples for the ecosystem-level impacts of human detritus that affects the environment directly as a pollutant, while also indirectly mediating ecological responses from feral and wild species (Barua & Sinha, 2022; Kumar et al., 2019). The reconfiguration of the regional biosphere by poultry and other livestock in tropical urban areas such as Delhi have created an ecological formation that links society and biota across multiple spatiotemporal scales (e.g., see Kumar, Gupta, et al., 2018, 2019). In most tropical cities, increasing solid waste that is disposed of informally, creates enormous foraging opportunities for urban fauna (Gutberlet, 2017). These steps involved in creation and culmination of organic biomass from waste highlight the need to better understand how animal husbandry activities in developing regions like South Asia intersect with socio-cultural, economic, and ecological processes. This case study at *Ghazipur* illustrated how animal-detritus has regional impacts on resident urban fauna, but also global effects on migratory avifauna (see Barua & Sinha, 2022). The largest landfill in New Delhi before the establishment of *Ghazipur* landfill in 1982 was located at *Indraprastha*, next to the National Zoological Park. Malhotra (2012) estimated that there were more than 40,000 Black-eared kites roosting in and around the zoo, unaware of Kumar et al.’s, (2020) study on the communal roosting habits of migratory Black-eared kites in Delhi. Since 1970s, the nesting density at Delhi’s Zoo has quadrupled (∼100 nests/km^2^: Kumar et al., 2019), and the roosting Black-eared kites’ numbers are usually less than 2000. It is also worth noting that even after the return of larger migratory kites, the number of resident breeding birds remained unchanged in *Ghazipur*, which highlights the variation in the significance of the site for the birds from two races. This finding supports the interesting fact discovered by Kumar et al., (2018) that the distance to a landfill does not influence the choice of breeding habitat by kite pairs in Delhi. These findings highlight major distinctions in the niches of resident vs. migratory kites, as well as breeding vs. non-breeding birds, suggesting potential areas of research on how life-history characteristics (migratory vs. resident) as well as individual life histories (young vs. old bird) shape and determine the exploitation of specific anthropogenic resources by urban commensals (Burnside et al., 2012; Hulme-Beaman et al., 2016; Seress & Liker, 2015). This highlights why poorly managed waste continues to have unpredictable and negative ecological consequences for human, animal, and environmental health across scales (Bettencourt et al., 2007; Grimm et al., 2008; Pickett et al., 2016).

The way that birds and other animals that eat dead things react to food from poultry farms, how infrastructure is built and changed in *Ghazipur* and SAK, and what people do - including how they relate to birds and the laws they make - all have an impact on how different types of plants and animals live together in cities (Barua, 2020). These communities are changing quickly and becoming more similar to each other (Fig. 5, 7) (Barua & Sinha, 2022; N. Kumar, Gupta, et al., 2018). It is critical to rethink the distribution of poultry waste and how it is influenced by urban infrastructure and socioeconomic factors. Furthermore, if we consider the estimates for GHG emissions, then the landfilled biomass from SAK constitutes anywhere between 4.23% to 26.81% of the annual methane (63.56 Gg/year) produced by landfilled biomass at *Ghazipur* in 2015, based on prior estimates by Ghosh et al., (2019). This shall enable recognition of its importance as a predictable foraging resource for both resident and migratory scavengers in cities like Delhi (see Fig. 6), which would have otherwise been overlooked. I discuss these further under specific headings (also see Kumar et al., 2019).

**Fig. 6.**
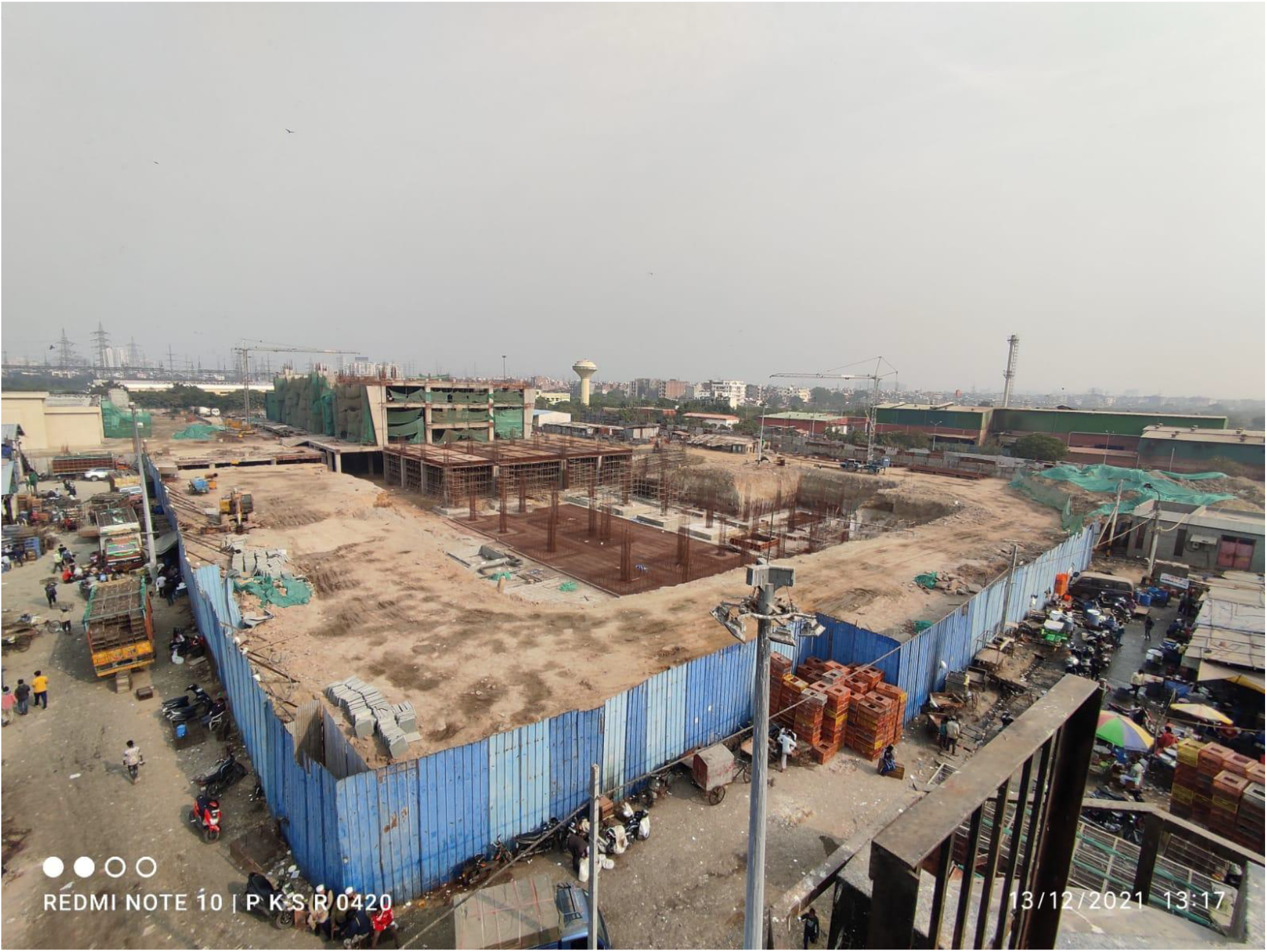
This aerial view shows the *Shaheed Ashfaqullah Khan* SAK, the largest poultry market in Asia. The poultry waste, produced by the daily trade of over 100,000 chickens, is an important food source for many opportunistic scavengers. In September 2018, the slaughter of poultry was halted following an order from the Delhi High Court, and a new processing facility has been under construction since then. The barricaded area that was previously used to collect poultry waste before it was loaded onto trucks (see Figures 2 and 3) also attracted a significant number of migratory kites. However, following the cessation of formal poultry slaughter (informal practices still ongoing), the number of kites has decreased to less than a third of the total congregated birds, resulting in a visible reduction to the kites’ photo captures (birds in flight), in comparison to the skyline in Fig. 1, 2 and 3.

### Implications for human health

The increasing prominence of broiler chicken in the human diet, ranking second only to pork, has resulted in a remarkable >400% increase in a broiler chicken’s body weight since 1950s (Bar-On et al., 2018; Bennett et al., 2018). This significant growth in poultry production has had direct dietary benefits and has created livelihood opportunities, particularly benefiting marginalised populations and women, thereby contributing to social and gender equity (Guèye, 2000; Jensen, n.d.). However, my findings reveal that inadequate disposal mechanisms for poultry waste in developing countries disproportionately impact urban poor communities, leading to caste and class-specific relationships with poultry waste, which can be classified as environmental racism (Gladkova, 2020; Pellow, 2004). Improper waste management practices have caused a multitude of ecological impacts, affecting the community of opportunistic scavengers in urban areas and resulting in direct conflicts with humans and other cohabiting species. A prime example of such conflicts is the burden of human-dog conflicts (bite), estimated to occur every two seconds in India by Menezes in 2008; these disproportionately affect the urban poor (Reporter, 2021). Issues of human-animal conflicts that are mediated by spatial dispersion of urban meat detritus highlight the global challenges of sharing living spaces with commensal species (Aiyedun & Olugasa, 2012; Gogtay et al., 2014; Hampson et al., 2015).

Existing interventions aimed at mitigating the implications of waste-mediated human-animal interfaces in shared spaces have proven inadequate. For instance, these interventions include selective neutering/translocation of stray dogs (Jackman & Rowan, 2007; Reese, 2005), managing monkey troops (Ganguly et al., 2018), implementing effective vaccination strategies against potential zoonotic diseases, culling of poultry and swine to prevent zoonotic influenza (Barua, 2020), and rehabilitating slums (Dupont & Gowda, 2020), and decommissioning landfills and abattoirs (Kumar et al., 2019). However, these approaches have not effectively addressed the ecological and behavioural relationships associated with waste and other forms of human food subsidies that enable certain animals to thrive in urban environments (S. Kumar et al., 2017; Schell et al., 2020; B. B. M. Wong & Candolin, 2015). A noteworthy organisational feature of PPC is its heavy reliance on precarious backyard farmers and workers that are urban immigrants. Based on the results of our semi-structured interviews, it is important to note how pandemic threats such as Avian Influenza and COVID-19 created a situation of virtual virulence in Delhi and beyond. Access to misinformation, such as WhatsApp forwards, devastated India’s poultry industry, which incurred losses in tune of $21 million/day in lockdown-I (Barua, 2020).

Telemetry data collected on kites (Kumar et al., 2020) and observations conducted at SAK have provided insights into how food subsidies influence the utilisation of airspace by both native and migratory bird species (Barua & Sinha, 2022). It is well-established that both native and exotic animal species thriving on poultry waste are associated with zoonotic concerns, thereby contributing to the perpetuation of antimicrobial resistance (AMR) pathways through the scavenger guild (Sharma et al., 2018). Notably, the interactions between rodents and kites over poultry and fish slaughter detritus are of particular concern, as they contribute to the imminent burden of AMR, which is predicted to experience the greatest growth in South Asia (Becker et al., 2018; Hassell et al., 2019; Khan et al., 2020; Woolhouse et al., 2015). In the Figure number 5, I illustrated the local and global impacts of the dominance of broiler chicken. These impacts, however, are often overlooked by the public health experts. It is important to note that socio-economic status within the Indian subcontinent plays a significant role in determining which parts of the chicken are consumed. Relatively wealthier individuals tend to discard the tarsii, skin, head and entrails, while in China, these parts are incorporated into highly regarded cuisines that are commonly consumed across caste and class lines (*pers. obs*). Understanding these dynamics is crucial for addressing the issue of AMR, taking into account the caste and class-specific consumption patterns associated with specific parts of poultry fowls in India (see Thieme, 2013, page. 20, for the contribution of each stage of the slaughter process to bacterial contamination).

The change in avian scavenger communities in South Asia following the decline of vulture populations is one aspect that has received little attention (Prakash et al., 2003). The rapid growth of the aviation industry in recent decades has resulted in an increase in the number of “aviation mega-cities” globally, projected to reach 91 by 2033, accompanied by an approximately 5% annual growth in air-passenger traffic over the next 20 years (Airbus, 2014). This expansion in air traffic, coupled with the preparations about utilisation of manned and unmanned drones for goods delivery and other purposes, creates a scenario where opportunistic birds and aircraft come into conflict within limited airspace. According to estimates provided by the Indian Air Force, birdstrikes affect approximately 1500 military and civilian aircraft annually in the region, leading to substantial financial losses and endangering human lives (IAF, *pers. comms.*). Among birds, avian scavengers, particularly Black Kites (*Milvus migrans*), pose the greatest threat in terms of birdstrike hazards, accounting for over 62% of all cases causing aircraft damage (IAF, *pers. comms.*). These conflicts would be less frequent if effective management strategies were employed for the disposal of poultry and other animal waste, highlighting the urgent need for novel approaches to wildlife management and conservation (Ditchkoff et al., 2006).

### Implications for animal health

Waste plays a significant role in shaping human-animal relationships, as it is often utilised as a direct and indirect food source for livestock and by opportunistic scavengers (Cavé, 2014; Kumar et al., 2019; Kumar et al., 2017; Sorathiya et al., 2014). In the context of Delhi, the availability of poultry waste is influenced by urban food habits, planning, and human beliefs and practices, with infrastructure playing a mediating role in determining access (Kumar et al., 2019). Through this research, I have revealed that the impact of poultry detritus on trophic dynamics exhibits high variability, dependent on the interplay between urban configuration and species-specific tolerance towards human proximity (see Kumar et al., 2019). For instance, the successful colonisation of detritus units by thousands of migratory kites in and around *Ghazipur* is contingent upon broader social perceptions of non-human life. Animals access anthropogenic resources such as poultry detritus, through species-specific movement patterns in response to built-up features like roads and overhead transmission wires they must negotiate (Criffield et al., 2018). The impacts of poultry detritus on trophic dynamics exhibit a high degree of geographic variability, which is contingent upon the spatially explicit coupling of urban configuration and species-specific tolerance towards human proximity (Kumar et al., 2019). The relationship between humans and animals is affected by a variety of factors, including the environment in which they live (extrinsic factors), the culture of the people involved, and the animals’ own characteristics (intrinsic factors). These factors can influence whether interactions between humans and animals are friendly, neutral, or hostile, and can also affect the way animals interact with each other (Sumasgutner et al., 2021). For instance, the wintering of thousands of Black-eared kites in and around the *Ghazipur* area is not only dependent on people’s social perceptions of non-human life in general, but also on the correlation between extrinsic urban habitat features within heterogeneously developed urban pockets and cultural geographies, combined with the intrinsic demographic factors of the animals (Kumar et al., 2019).

Furthermore, urbanisation, which offers predictable food subsidies, has likely impacted threatened migratory facultative scavengers such as Steppe Eagles and Eurasian Griffon vultures that congregate at landfills and carcass dumping sites (*pers obs.*). Despite its impact on biodiversity, urban expansion is an inevitable outcome of development. The rise of cities along the migratory flyways not only offers novel anthropogenic-opportunities to avian scavengers, but also multiple threats from overhead transmission wires, wind energy farms, buildings, artificial-lights at night, etc. (A. L. Drewitt & Langston, 2008; E. J. A. Drewitt et al., 2021; Horton et al., 2019; N. Kumar et al., 2020; Nilsson et al., 2021). But development is still a work in progress for vast tropical landscapes. It offers opportunities to allow incorporation of wireless technology in planning ‘green’ and ‘smart’ infrastructure in the Global South that can support long-term coexistence of birdlife and people linked through ecosystem services, e.g., safe disposal of human-refuse by opportunistic avian migrants and scavengers, and their cultural reverence (Natuhara, 2018). In tropical countries, migratory birds and humans often coexist in urban settings for long periods of time. This is because the birds interact with anthropogenic resources that are shaped by sociocultural and economic factors. These dynamic nature-culture relationships also lead to unique model systems for ecological and ethno-ornithological studies (Colin A. Galbraith, Tim Jones, Jeff Kirby and Taej Mundkur, 2014; Mikula et al., 2023; Newton, 2010). The results of this study and observations from carcass dumping sites like *Jorbeer* (Rajasthan) (Khatri, 2013) and several areas in Uttar Pradesh (Jha, 2015) suggest that migratory avian scavengers have the potential to have a significant impact on conservation and public awareness of science. For instance, they can lead to revenue-generating models from bird-watching tourism and small industries (e.g., bovine-leather and bone-meal fertilisers near carcass-dumping grounds) (MOEF&CC, 2020; Sinha, 1986). Regarding ethnic-philosophy that is well known to support and care non-human lives in South Asia, Indian-scriptures mention “non-violence as the ultimate duty” 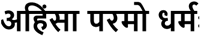:/ *Ahimsa-Parmo-Dharma*) and teach to “respect and honour guests as God” (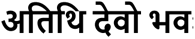:/*Atithi-Devo-Bhava*) (Dave, 2005). Currently, these ethnic principles are threatened by the simultaneous loss of biodiversity and cultures in urbanising tropical cities (N. Kumar, Gupta, et al., 2018).

> It is worth highlighting an incident that occurred in April 2017, which exemplifies the phenomenon of “empathic reinforcement” between daily-waged immigrant butchers/cleaners and kites at SAK in Delhi (*Fig. 3-6*). Upon observing the migration routes of kites between Delhi and Central Asia on Google Maps, members of the community demonstrated compassion by providing care to 15-20 Black-eared kites that were in a moribund state after a dust-storm. They offered these birds food, water, and temporary shelter. This response stemmed from the innate love for living creatures, often referred to as biophilia, which finds expression in religious beliefs prevalent in South Asia (Pinault, 2008; Taneja, 2015). Both Muslims and Hindus hold reverence for kites, albeit with different perspectives. For Muslims, kites are regarded as “winged emissaries” that carry away sins, worries, or prayers-messages, symbolized by the act of offering meat to these birds during ritual feeding near mosques. On the other hand, Hindus perceive interconnectedness among all life forms, viewing animal species as holy beings or even solutions to astrological problems, thus displaying tolerance towards usual inconvenience caused by aggressive animal parents, e.g., kites, dogs, and macaques (Kumar et al., 2019; Pinault, 2008; Taneja, 2015).

Delhi residents are sympathetic towards aggressive co-inhabiting animals because of the mutualism derived from animals’ parenting behaviour and from the realisation that urbanisation has destroyed and degraded the natural habitats of wildlife. Interviews revealed that residents were well aware of the useful “scavenging ecosystem services” provided to their neighbourhood by kites and other animals which consume poultry and other detritus (see Kumar at al., 2019). Therefore, unintended ecological consequences of landfill flattening can manifest in various ways, including the hazards posed to bird populations due to increased birdstrike incidents, movement, migration and/or dispersal of animals in search of new resources, and the potential contamination of soil and water resources used by various life forms (Ahluwalia & Patel, 2018; Plaza & Lambertucci, 2017; Poore & Nemecek, 2018). While it is beyond the scope of this manuscript, by thoroughly considering and addressing these factors, we can minimise negative ecological impacts on humans/animals and ensure the sustainable management of waste materials during landfill flattening. In summary, while the current political discussions surrounding landfill flattening in Delhi highlight a progressive approach to waste management, it is imperative to conduct a comprehensive evaluation of the ecological implications and challenges associated with these initiatives. By carefully considering the organic composition of waste materials and implementing appropriate mitigation strategies, we can mitigate potential environmental risks and contribute to sustainable waste management practices in the context of a tropical megacity (Ahluwalia & Patel, 2018; Pellow, 2004; Shahmoradi, 2013).

### Implications for environmental health

Six percent of global greenhouse emissions come from food losses and waste (Poore & Nemecek, 2018). This study found that a large amount of poultry waste is left to decompose in open spaces or drains because of the lack of proper disposal methods, especially during the threat of avian influenza (Gržinić et al., 2023). The traditional approach of relying on scavenging services provided by commensal animals that works on small village-town scales (see Kumar et al., 2019) has been found to create conflicts and contribute to the spread of diseases (Nyhus, 2016; Schwarzenbach et al., 2010; Scoones, 2016; Vergara & Tchobanoglous, 2012). Although this paper does not directly address the issue, large-scale poultry production raises environmental concerns, such as the high carbon footprint of chicken feed and the high nitrogen content of chicken manure (Gržinić et al., 2023). Suggestions have been made to address the environmental impact of global poultry dominance, including the production of biodiesel, biogas, organic manure, and compost (Faraji Mahyari et al., 2021; Gerber et al., 2008; Wang et al., 2014). In cities like Delhi, the problem of poorly managed poultry waste is not only spatial but also seasonal. The absence of Black-eared kites, which primarily scavenge on such remains at *Ghazipur*, during the six summer months increases the risk of disease transmission from decaying carrion, especially given the proliferation of insect vectors during warm and humid weather conditions (Krystosik et al., 2020). There lies a significant global challenge in optimising these tropical urban spaces to support PPC activities while minimising the health risks associated with cohabiting with animals that have adapted to urban environments (Council, 2001).

Human socio-cultural factors play a significant role in determining the settlement patterns of both people and animals around poultry waste, which can lead to physical conflicts between humans and wildlife or the transmission of zoonotic diseases (Krystosik et al., 2020; N. Kumar et al., 2020; N. Kumar, Jhala, et al., 2019; Olson et al., 2016; Onen & Bassey, 2017). Migratory birds, such as waterfowl, are known carriers of avian influenza. They have been observed roosting in close proximity to each other, posing a risk of spreading infectious diseases to poultry, livestock, humans, and even endangered wildlife (Bobiec et al., 2021; Bradley et al., 2020; Grace et al., 2012; N. Kumar et al., 2020; Xu et al., 2016). The environmental burden associated with poultry waste is influenced by the geography of human religious practices, hygiene practices, and poverty. More often than not, in tropical cities that have heterogeneously developed pockets, gentrification drives result in concentrated waste accumulation at the outskirts of the city. In addition, *Ghazipur* in Delhi has become a prime example of how a lack of foresight about systemic linkages in health and hygiene, and informal to formal eco-commercial linkages can create a variety of problems affecting humans and other living things (Ferronato & Torretta, 2019; Gutberlet, 2017; S. Kumar et al., 2017; Talyan et al., 2008). Consequently, the process of landfill flattening, and waste repurposing should be carried out with caution to protect the environmental and social well-being of the surrounding ecological communities supported by and centred on humans. Appropriate measures should be taken to address potential issues, such as the release of harmful GHG and pathogens from decomposing waste, which can cause air, land, and water pollution (Krystosik et al., 2020).

The predictable distribution of food waste from humans has been shown to affect the behaviour of urban animals such as kites (see Kumar et al., 2018, 2019). Kites tolerated humans at close range when feeding on carrion but will attack if provoked near the nests (Kumar, Jhala, et al., 2019; N. Kumar, Qureshi, et al., 2018). This was not observed for the birds from the migratory *lineatus* race that fled on approach, much like the behaviour of the resident *govinda* kites when not breeding (*unpiblished data*). Such “coupled-systems” in which humans and non-human species interact and shape each other’s socio-ecological space through repeated interactions highlights the impact of urbanisation on ethno-zoological domains in megacities (see Kumar et al., 2019). The age-old human-scavenger relationships highlighted by this study for a megacity are facing a growing threat from the progressive lack of appreciation by younger generations of the value of intangible benefits provided by wildlife to humans (Curtin & Kragh, 2014). It is critical to educate younger generations about the importance of wildlife so that these relationships can be maintained. This human-nature disconnect has alienated people through the “extinction of experience,” with simultaneous consequences for human, animal, and environmental health (Kumar et al., 2019). Addressing this issue necessitates focused research and public policy on documenting and promoting biocultural awareness and research about the human-nature disconnect and develop public policies to address it (Soga & Gaston, 2016).

## Conclusion

This study has provided valuable insights into the management of poultry detritus in Delhi, highlighting its significant implications for environmental and public health (Dunn et al., 2006; Gržinić et al., 2023). The traditional practice of benefitting from scavenging services of animals that help dispose of waste at the scales of villages and towns is undergoing rapid change, often leading to human-animal conflicts that contribute to the spread of zoonotic diseases. The geographic dispersion of anthropogenic food subsidies in the region, which is influenced by human socio-cultural factors such as religion, hygiene, and poverty, plays a significant role in determining the distribution of environmental burdens. The interconnected relationship between humans and non-human taxa, as observed in this study, emphasises the impact of urbanisation on ethno-zoological domains in megacities. Understanding and addressing this coupled-system, where humans and wildlife mutually shape each other’s social-ecological space through repeated interactions, is essential in mitigating conflicts and promoting harmonious coexistence. It is anticipated that conflicts arising from close exposure to humans will increase as growing economies and urban populations worldwide attract predatory vertebrates with similar subsidies. To effectively manage urban wildlife and minimise conflicts, it is crucial to incorporate ethno-zoological insights into ecological considerations. In addition, relying solely on waste-to-value chains that use human waste as food subsidies for non-human species may not be ecologically sustainable. It is important to carefully examine its impacts on human, animal, and environmental health, particularly the release of GHG, zoonosis, and antimicrobial resistance.

If food waste were a country, its capacity to produce geen house gases would globally rank third, after the USA and China (FAO, 2023). Therefore, as tropical regions like India prepare to urbanise and accommodate more than 2.5 billion citizens in the coming decades, it becomes imperative to optimise urban spaces for humans, solid waste management, and human-animal coexistence. A holistic approach that considers ecological and socio-cultural factors is needed for effective management of waste from animal processing for food, e.g., poultry. In summary, this study underscores the importance of adopting a comprehensive approach to poultry detritus management, considering the ecological and socio-cultural dynamics that shape the urban landscape. As cities expand and human-wildlife interactions become more prevalent, promoting biocultural awareness, and integrating ecological insights can foster the development of sustainable and “waste wise” urban ecosystems (Becker et al., 2015, 2018).

## Acknowledgement

I would like to thank the following people and organisations for their help in making this work possible: The **PAWS-Web Network** (www.thinkPAWS.org) for their workshops and discussions. The Felix Scholarship Trust, India Oxford Initiative and grants from the University of Oxford, the Raptor Research and Conservation Foundation, Mumbai, and the Government of India’s Ministry of Environment Forest & Climate Change for funding. Delhi’s Police, Department of Forest & Wildlife, National Zoological Park, Municipal Corporations, and civic bodies for giving permits and logistic support. Yadvendradev V Jhala, Qamar Qureshi, Andrew Gosler and Fabrizio Sergio for supervising the Black Kite Project/my masters and doctoral dissertaions. Urvi, Nikita, Amee, Yukti, Laxmi, Prince, Jugnu, Zehen, Sourabh, Nitish, Aparajita, Prashant, Harsha, Poonam, Abhinandan, Navneet and Gunjesh (and more than 200 volunteers/interns who worked with the Black Kite Project) for fieldwork during 2012 -23. Delhi citizens and officials at the Wildlife Institute of India for their support. I am grateful for their contributions and support. Comments from Ujjwal Kumar and Apoorva Kulkarni improved previous drafts.

## References

Ahluwalia, I. J., & Patel, U. (2018). Solid waste management in India: An assessment of resource recovery and environmental impact.

Airbus. (2014). Airbus: Emerging markets and urbanization driving air passenger traffic growth; up 4.7% annually for next 20 years. Green Car Congress. https://www.greencarcongress.com/2014/09/20140925-airbus.html

Aiyedun, J. O., & Olugasa, B. O. (2012). Identification and analysis of dog use, management practices and implications for rabies control in Ilorin, Nigeria. Sokoto Journal of Veterinary Sciences, 10(2), Article 2. https://doi.org/10.4314/sokjvs.v10i2.1

Bar-On, Y. M., Phillips, R., & Milo, R. (2018). The biomass distribution on Earth. Proceedings of the National Academy of Sciences, 115(25), 6506–6511. https://doi.org/10.1073/pnas.1711842115

Barua, M. (2020, April 17). Virtual Virulence and Metabolic Life. Society for Cultural Anthropology. https://culanth.org/fieldsights/virtual-virulence-and-metabolic-life

Barua, M., & Sinha, A. (2022). Cultivated, feral, wild: The urban as an ecological formation. Urban Geography, 0(0), 1–22. https://doi.org/10.1080/02723638.2022.2055924

Becker, D. J., Hall, R. J., Forbes, K. M., Plowright, R. K., & Altizer, S. (2018). Anthropogenic resource subsidies and host–parasite dynamics in wildlife. Philosophical Transactions of the Royal Society B: Biological Sciences, 373(1745), 20170086. https://doi.org/10.1098/rstb.2017.0086

Becker, D. J., Streicker, D. G., & Altizer, S. (2015). Linking anthropogenic resources to wildlife– pathogen dynamics: A review and meta-analysis. Ecology Letters, 18(5), 483–495. https://doi.org/10.1111/ele.12428

Bennett, C. E., Thomas, R., Williams, M., Zalasiewicz, J., Edgeworth, M., Miller, H., Coles, B., Foster, A., Burton, E. J., & Marume, U. (2018). The broiler chicken as a signal of a human reconfigured biosphere. Royal Society Open Science, 5(12), 180325. https://doi.org/10.1098/rsos.180325

Bettencourt, L. M. A., Lobo, J., Helbing, D., Kühnert, C., & West, G. B. (2007). Growth, innovation, scaling, and the pace of life in cities. Proceedings of the National Academy of Sciences, 104(17), 7301–7306. https://doi.org/10.1073/pnas.0610172104

Bhagat, R. B., & Mohanty, S. (2009). Emerging Pattern of Urbanization and the Contribution of Migration in Urban Growth in India. Asian Population Studies, 5(1), 5–20. https://doi.org/10.1080/17441730902790024

BJP Manifesto. (2022, November 25). BJP releases “Sankalp Patra” for Delhi municipal polls; online services. Deccan Chronicle. https://www.deccanchronicle.com/nation/politics/251122/bjp-releases-sankalp-patra-for-delhi-municipal-polls-online-service.html

Bobiec, A., Paderewski, J., & Gajdek, A. (2021). Urbanisation and globalised environmental discourse do not help to protect the bio-cultural legacy of rural landscapes. Landscape and Urban Planning, 208, 104038. https://doi.org/10.1016/j.landurbplan.2021.104038

Bourdeau-Lepage, L., & Huriot, J.-M. (2007). Megacities without global functions. Belgeo. Revue Belge de Géographie, 1, Article 1. https://doi.org/10.4000/belgeo.11675

Bradley, A., Mennie, N., Bibby, P. A., & Cassaday, H. J. (2020). Some animals are more equal than others: Validation of a new scale to measure how attitudes to animals depend on species and human purpose of use. PLOS ONE, 15(1), e0227948. https://doi.org/10.1371/journal.pone.0227948

Burnside, W. R., Brown, J. H., Burger, O., Hamilton, M. J., Moses, M., & Bettencourt, L. M. A. (2012). Human macroecology: Linking pattern and process in big-picture human ecology. Biological Reviews, 87(1), 194–208. https://doi.org/10.1111/j.1469-185X.2011.00192.x

Cavé, J. (2014). Who owns urban waste? Appropriation conflicts in emerging countries. Waste Management & Research, 32(9), 813–821. https://doi.org/10.1177/0734242X14540978

*Census* 2011 India. (2011). https://www.census2011.co.in/

Champion, H. G., & Seth, S. K. (1968). A revised survey of the forest types of India. Manager of Publications. https://dds.crl.edu/crldelivery/23005

Chatterjee, R. N., & Rajkumar, U. (2015). AN OVERVIEW OF POULTRY PRODUCTION IN INDIA.

Colin A. Galbraith, Tim Jones, Jeff Kirby and Taej Mundkur. (2014). A Review of Migratory Bird Flyways and Priorities for Management: CMS Technical Series Publication *No*. 27. UNEP / CMS Secretariat, Bonn, Germany.

Council, N. R. (2001). Under the Weather: Climate, Ecosystems, and Infectious Disease. The National Academies Press. https://doi.org/10.17226/10025

Criffield, M., van de Kerk, M., Leone, E., Cunningham, M. W., Lotz, M., Oli, M. K., & Onorato, D. P. (2018). Assessing impacts of intrinsic and extrinsic factors on Florida panther movements. Journal of Mammalogy, 99(3), 702–712. https://doi.org/10.1093/jmammal/gyy025

Curtin, S., & Kragh, G. (2014). Wildlife Tourism: Reconnecting People with Nature. Human Dimensions of Wildlife, 19(6), 545–554. https://doi.org/10.1080/10871209.2014.921957

Dave, K. N. (2005). Birds in Sanskrit Literature: With 107 Bird Illustrations. Motilal Banarsidass Publishe.

Devi, S. M., Balachandar, V., Lee, S. I., & Kim, I. H. (2014). An Outline of Meat Consumption in the Indian Population—A Pilot Review. Korean Journal for Food Science of Animal Resources, 34(4), 507–515. https://doi.org/10.5851/kosfa.2014.34.4.507

Ditchkoff, S. S., Saalfeld, S. T., & Gibson, C. J. (2006). Animal behavior in urban ecosystems: Modifications due to human-induced stress. Urban Ecosystems, 9(1), 5–12. https://doi.org/10.1007/s11252-006-3262-3

Dopelt, K., Radon, P., & Davidovitch, N. (2019). Environmental Effects of the Livestock Industry: The Relationship between Knowledge, Attitudes, and Behavior among Students in Israel. International Journal of Environmental Research and Public Health, 16(8), 1359. https://doi.org/10.3390/ijerph16081359

Doron, A. (2021). Stench and sensibilities: On living with waste, animals and microbes in India. The Australian Journal of Anthropology, 32(S1), 23–41. https://doi.org/10.1111/taja.12380

Drewitt, A. L., & Langston, R. H. W. (2008). Collision Effects of Wind-power Generators and Other Obstacles on Birds. Annals of the New York Academy of Sciences, 1134(1), 233–266. https://doi.org/10.1196/annals.1439.015

Drewitt, E. J. A., Cuthill, I. C., Sutton, L. J., Smith, H. R., Loram, S. W., & Thomas, R. J. (2021). Northerly dispersal trends in a lowland population of Peregrines Falco peregrinus in southwest England. Ringing & Migration, 36(2), 105–115. https://doi.org/10.1080/03078698.2022.2150783

Dunn, R. R., Gavin, M. C., Sanchez, M. C., & Solomon, J. N. (2006). The Pigeon Paradox: Dependence of Global Conservation on Urban Nature. Conservation Biology, 20(6), 1814–1816. https://doi.org/10.1111/j.1523-1739.2006.00533.x

Dupont, V., & Gowda, M. M. S. (2020). Slum-free city planning versus durable slums. Insights from Delhi, India. International Journal of Urban Sustainable Development, 12(1), 34–51. https://doi.org/10.1080/19463138.2019.1666850

Esmail, N., Wintle, B. C., t Sas-Rolfes, M., Athanas, A., Beale, C. M., Bending, Z., Dai, R., Fabinyi, M., Gluszek, S., Haenlein, C., Harrington, L. A., Hinsley, A., Kariuki, K., Lam, J., Markus, M., Paudel, K., Shukhova, S., Sutherland, W. J., Verissimo, D., … Milner-Gulland, E. J. (2020). Emerging illegal wildlife trade issues: A global horizon scan. Conservation Letters, 13(4), e12715. https://doi.org/10.1111/conl.12715

FAO. (2019). World Livestock: Transforming the livestock sector through the Sustainable Development Goals. FAO. https://doi.org/10.4060/ca1201en

FAO. (2023). Food Wastage Footprint & Climate Change. http://www.fao.org/nr/sustainability/food-loss-and-waste

FAO 2022. (n.d.). Products and processing | Gateway to poultry production and products | Food and Agriculture Organization of the United Nations. Retrieved April 16, 2022, from https://www.fao.org/poultry-production-products/products-processing/en/

Faraji Mahyari, Z., Khorasanizadeh, Z., Khanali, M., & Faraji Mahyari, K. (2021). Biodiesel production from slaughter wastes of broiler chicken: A potential survey in Iran. SN Applied Sciences, 3(1), 57. https://doi.org/10.1007/s42452-020-04045-7

Ferronato, N., & Torretta, V. (2019). Waste Mismanagement in Developing Countries: A Review of Global Issues. International Journal of Environmental Research and Public Health, 16(6), 1060. https://doi.org/10.3390/ijerph16061060

Gangoso, L., Agudo, R., Anadón, J. D., de la Riva, M., Suleyman, A. S., Porter, R., & Donázar, J. A. (2013). Reinventing mutualism between humans and wild fauna: Insights from vultures as ecosystem services providers. Conservation Letters, 6(3), 172–179. https://doi.org/10.1111/j.1755-263X.2012.00289.x

Ganguly, I., Chauhan, N. P. s, & Verma, P. (2018). Assessment of Human-Macaque Conflict and Possible Mitigation Strategies in and Around Asola-Bhatti Wildlife Sanctuary, Delhi NCR.

Gerber, P. J., Opio, C. I., & Steinfeld, H. (2008). Poultry production and the environment – a review.

Ghosh, P., Shah, G., Chandra, R., Sahota, S., Kumar, H., Vijay, V. K., & Thakur, I. S. (2019). Assessment of methane emissions and energy recovery potential from the municipal solid waste landfills of Delhi, India. Bioresource Technology, 272, 611–615. https://doi.org/10.1016/j.biortech.2018.10.069

Gilbert, M., Conchedda, G., Boeckel, T. P. V., Cinardi, G., Linard, C., Nicolas, G., Thanapongtharm, W., D’Aietti, L., Wint, W., Newman, S. H., & Robinson, T. P. (2015). Income Disparities and the Global Distribution of Intensively Farmed Chicken and Pigs. PLOS ONE, 10(7), e0133381. https://doi.org/10.1371/journal.pone.0133381

Gilbert, N. I., Correia, R. A., Silva, J. P., Pacheco, C., Catry, I., Atkinson, P. W., Gill, J. A., & Franco, A. M. A. (2016). Are white storks addicted to junk food? Impacts of landfill use on the movement and behaviour of resident white storks (Ciconia ciconia) from a partially migratory population. Movement Ecology, 4(1), 7. https://doi.org/10.1186/s40462-016-0070-0

Gil-Fernández, M., Harcourt, R., Newsome, T., Towerton, A., & Carthey, A. (2020). Adaptations of the red fox (Vulpes vulpes) to urban environments in Sydney, Australia. Journal of Urban Ecology, 6(1), juaa009. https://doi.org/10.1093/jue/juaa009

Gladkova, E. (2020). Farming Intensification and Environmental Justice in Northern Ireland. Critical Criminology, 28(3), 445–461. https://doi.org/10.1007/s10612-020-09488-3

Gogtay, N. J., Nagpal, A., Mallad, A., Patel, K., Stimpson, S. J., Belur, A., & Thatte, U. M. (2014). Demographics of animal bite victims & management practices in a tertiary care institute in Mumbai, Maharashtra, India. The Indian Journal of Medical Research, 139(3), 459–462.

GoI. (2019). 20th Livestock Census-2019: All India Report. https://ruralindiaonline.org/en/library/resource/20th-livestock-census-2019-all-india-report/

GoI. (2022). Animal Husbandry Statistics (AHS) | Department of Animal Husbandry & Dairying. https://dahd.nic.in/animal-husbandry-statistics

Grace, D., Mutua, F., Ochungo, P., Kruska, R. L., Jones, K., Brierley, L., Lapar, M. L., Said, M. Y., Herrero, M. T., Phuc, P. M., Thao, N. B., Akuku, I., & Ogutu, F. (2012). Mapping of poverty and likely zoonoses hotspots [Report]. International Livestock Research Institute. https://cgspace.cgiar.org/handle/10568/21161

Griffin, A. S., Netto, K., & Peneaux, C. (2017). Neophilia, innovation and learning in an urbanized world: A critical evaluation of mixed findings. Current Opinion in Behavioral Sciences, 16, 15–22. https://doi.org/10.1016/j.cobeha.2017.01.004

Griffin, L. L., Haigh, A., Amin, B., Faull, J., Norman, A., & Ciuti, S. (2022). Artificial selection in human-wildlife feeding interactions. Journal of Animal Ecology, 91(9), 1892–1905. https://doi.org/10.1111/1365-2656.13771

Grimm, N. B., Faeth, S. H., Golubiewski, N. E., Redman, C. L., Wu, J., Bai, X., & Briggs, J. M. (2008). Global Change and the Ecology of Cities. Science, 319(5864), 756–760. https://doi.org/10.1126/science.1150195

Gržinić, G., Piotrowicz-Cieślak, A., Klimkowicz-Pawlas, A., Górny, R. L., Ławniczek-Wałczyk, A., Piechowicz, L., Olkowska, E., Potrykus, M., Tankiewicz, M., Krupka, M., Siebielec, G., & Wolska, L. (2023). Intensive poultry farming: A review of the impact on the environment and human health. Science of The Total Environment, 858, 160014. https://doi.org/10.1016/j.scitotenv.2022.160014

Guèye, E. F. (2000). The Role of Family Poultry in Poverty Alleviation, Food Security and the Promotion of Gender Equality in Rural Africa. Outlook on Agriculture, 29(2), 129–136. https://doi.org/10.5367/000000000101293130

Gutberlet, J. (2017). Waste in the City: Challenges and Opportunities for Urban Agglomerations. In Urban Agglomeration. IntechOpen. https://doi.org/10.5772/intechopen.72047

Hampson, K., Coudeville, L., Lembo, T., Sambo, M., Kieffer, A., Attlan, M., Barrat, J., Blanton, J. D., Briggs, D. J., Cleaveland, S., Costa, P., Freuling, C. M., Hiby, E., Knopf, L., Leanes, F., Meslin, F.-X., Metlin, A., Miranda, M. E., Müller, T., … Prevention, on behalf of the G. A. for R. C. P. for R. (2015). Estimating the Global Burden of Endemic Canine Rabies. PLOS Neglected Tropical Diseases, 9(4), e0003709. https://doi.org/10.1371/journal.pntd.0003709

Hassell, J. M., Ward, M. J., Muloi, D., Bettridge, J. M., Robinson, T. P., Kariuki, S., Ogendo, A., Kiiru, J., Imboma, T., Kang’ethe, E. K., Öghren, E. M., Williams, N. J., Begon, M., Woolhouse, M. E. J., & Fèvre, E. M. (2019). Clinically relevant antimicrobial resistance at the wildlife–livestock– human interface in Nairobi: An epidemiological study. The Lancet Planetary Health, 3(6), e259– e269. https://doi.org/10.1016/S2542-5196(19)30083-X

Hendrix, P. F., Parmelee, R. W., Crossley, D. A., Coleman, D. C., Odum, E. P., & Groffman, P. M. (1986). Detritus Food Webs in Conventional and No-Tillage Agroecosystems. BioScience, 36(6), 374–380. https://doi.org/10.2307/1310259

Horton, K. G., Nilsson, C., Van Doren, B. M., La Sorte, F. A., Dokter, A. M., & Farnsworth, A. (2019). Bright lights in the big cities: Migratory birds’ exposure to artificial light. Frontiers in Ecology and the Environment, 17(4), 209–214. https://doi.org/10.1002/fee.2029

Hulme-Beaman, A., Dobney, K., Cucchi, T., & Searle, J. B. (2016). An ecological and evolutionary framework for commensalism in anthropogenic environments. Trends in Ecology & Evolution, 31(8), Article 8.

IMD. (2022). Indian Metrological Department: New Delhi. https://mausam.imd.gov.in/Delhi/

Jackman, J., & Rowan, A. (2007). Free-Roaming Dogs in Developing Countries: The Benefits of Capture, Neuter, and Return Programs. State of the Animals 2007. https://www.wellbeingintlstudiesrepository.org/sota_2007/10

Jensen, H. A. (n.d.). Poultry as a Tool in Poverty Eradication and Promotion of Gender Equality. Retrieved May 3, 2022, from https://www.fao.org/3/ac154e/AC154E02.htm

Jha, K. K. (2015). Distribution of vultures in Uttar Pradesh, India. Journal of Threatened Taxa, 7(1), Article 1. https://doi.org/10.11609/JoTT.o3319.6750-63

Karwowska, M., Łaba, S., & Szczepański, K. (2021). Food Loss and Waste in Meat Sector—Why the Consumption Stage Generates the Most Losses? Sustainability, 13(11), Article 11. https://doi.org/10.3390/su13116227

Katlam, G., Prasad, S., Aggarwal, M., & Kumar, R. (2018). Trash on the menu. Current Science, 115(12), 2322–2326. JSTOR.

Khan, S. A., Imtiaz, M. A., Sayeed, Md. A., Shaikat, A. H., & Hassan, M. M. (2020). Antimicrobial resistance pattern in domestic animal—Wildlife—Environmental niche via the food chain to humans with a Bangladesh perspective; a systematic review. BMC Veterinary Research, 16(1), 302. https://doi.org/10.1186/s12917-020-02519-9

Khatri, P. C. (2013). HOME RANGE USE OF WINTER MIGRATORY VULTURES IN AND AROUND JORBEER, BIKANER (RAJASTHAN) INDIA.

Krystosik, A., Njoroge, G., Odhiambo, L., Forsyth, J. E., Mutuku, F., & LaBeaud, A. D. (2020). Solid Wastes Provide Breeding Sites, Burrows, and Food for Biological Disease Vectors, and Urban Zoonotic Reservoirs: A Call to Action for Solutions-Based Research. Frontiers in Public Health, 7, 405. https://doi.org/10.3389/fpubh.2019.00405

Kumar, N. (2019). Urban Waste and the Human–Animal Interface in Delhi | Economic and Political Weekly. https://www.epw.in/journal/2019/47/review-urban-affairs/urban-waste--human-animal-interface-delhi.html

Kumar, N. (2013). A Study of Resource Selection by Black Kites Milvus migrans in the Urban Landscape of National Capital Region, India. M.Sc. Thesis. Wildlife Institute of India. https://www.raptors.org.in/dissertations/

Kumar, N., Gupta, U., Jhala, Y. V., Qureshi, Q., Gosler, A. G., & Sergio, F. (2018). Habitat selection by an avian top predator in the tropical megacity of Delhi: Human activities and socio-religious practices as prey-facilitating tools. Urban Ecosystems, 21(2), 339–349. https://doi.org/10.1007/s11252-017-0716-8

Kumar, N., Gupta, U., Jhala, Y. V., Qureshi, Q., Gosler, A. G., & Sergio, F. (2020). GPS-telemetry unveils the regular high-elevation crossing of the Himalayas by a migratory raptor: Implications for definition of a “Central Asian Flyway.” Scientific Reports, 10(1), Article 1. https://doi.org/10.1038/s41598-020-72970-z

Kumar, N., Gupta, U., Malhotra, H., Jhala, Y. V., Qureshi, Q., Gosler, A. G., & Sergio, F. (2019). The population density of an urban raptor is inextricably tied to human cultural practices. Proceedings of the Royal Society B: Biological Sciences, 286(1900), 20182932. https://doi.org/10.1098/rspb.2018.2932

Kumar, N., Jhala, Y. V., Qureshi, Q., Gosler, A. G., & Sergio, F. (2019). Human-attacks by an urban raptor are tied to human subsidies and religious practices. Scientific Reports, 9(1), Article 1. https://doi.org/10.1038/s41598-019-38662-z

Kumar, N., Qureshi, Q., Jhala, Y. V., Gosler, A. G., & Sergio, F. (2018). Offspring defense by an urban raptor responds to human subsidies and ritual animal-feeding practices. PLOS ONE, 13(10), e0204549. https://doi.org/10.1371/journal.pone.0204549

Kumar, S., Smith, S. R., Fowler, G., Velis, C., Kumar, S. J., Arya, S., Rena, null, Kumar, R., & Cheeseman, C. (2017). Challenges and opportunities associated with waste management in India. Royal Society Open Science, 4(3), 160764. https://doi.org/10.1098/rsos.160764

Malhotra, D. A. K. (2012). Tiger of the Sky: Pariah Kite. SHILALEKH.

Marzluff, J. M. (2017). A decadal review of urban ornithology and a prospectus for the future. Ibis, 159(1), 1–13. https://doi.org/10.1111/ibi.12430

Menezes, R. (2008). Public health: Rabies in India. CMAJ : Canadian Medical Association Journal, 178(5), 564. https://doi.org/10.1503/cmaj.071488

Mikula, P., Tomášek, O., Romportl, D., Aikins, T. K., Avendaño, J. E., Braimoh-Azaki, B. D. A., Chaskda, A., Cresswell, W., Cunningham, S. J., Dale, S., Favoretto, G. R., Floyd, K. S., Glover, H., Grim, T., Henry, D. A. W., Holmern, T., Hromada, M., Iwajomo, S. B., Lilleyman, A., … Albrecht, T. (2023). Bird tolerance to humans in open tropical ecosystems. Nature Communications, 14(1), Article 1. https://doi.org/10.1038/s41467-023-37936-5

MOEF&CC. (2020). *Action Plan for Vulture Conservation in India 2020-*2025. https://save-vultures.org/action-plans/

Moleón, M., Sánchez-Zapata, J. A., Margalida, A., Carrete, M., Owen-Smith, N., & Donázar, J. A. (2014). Humans and Scavengers: The Evolution of Interactions and Ecosystem Services. BioScience, 64(5), 394–403. https://doi.org/10.1093/biosci/biu034

Mozhiarasi, V., & Natarajan, T. S. (2022). Slaughterhouse and poultry wastes: Management practices, feedstocks for renewable energy production, and recovery of value-added products. Biomass Conversion and Biorefinery. https://doi.org/10.1007/s13399-022-02352-0

MCRS Music. (2020). Murga Badnaam Huwa Corona Tere Liye. Retrieved July 12, 2023, from https://www.youtube.com/watch?v=UC38a_Pz_Og.

Mukherjee, N., Hugé, J., Sutherland, W. J., McNeill, J., Van Opstal, M., Dahdouh-Guebas, F., & Koedam, N. (2015). The Delphi technique in ecology and biological conservation: Applications and guidelines. Methods in Ecology and Evolution, 6(9), 1097–1109. https://doi.org/10.1111/2041-210X.12387

Natuhara, Y. (2018). Green infrastructure: Innovative use of indigenous ecosystems and knowledge. Landscape and Ecological Engineering, 14(2), 187–192. https://doi.org/10.1007/s11355-018-0357-y

Newton, I. (2010). The Migration Ecology of Birds. Elsevier.

Nilsson, C., La Sorte, F. A., Dokter, A., Horton, K., Van Doren, B. M., Kolodzinski, J. J., Shamoun-Baranes, J., & Farnsworth, A. (2021). Bird strikes at commercial airports explained by citizen science and weather radar data. Journal of Applied Ecology, 58(10), 2029–2039. https://doi.org/10.1111/1365-2664.13971

Njuki, J., Waithanji, E., Bagalwa, N., & Kariuki, J. (2013). Guidelines on integrating gender in livestock projects and programs. https://cgspace.cgiar.org/bitstream/handle/10568/33425/GenderInLivestock.pdf

Nyhus, P. J. (2016). Human–Wildlife Conflict and Coexistence. Annual Review of Environment and Resources, 41(1), 143–171. https://doi.org/10.1146/annurev-environ-110615-085634

Olson, Z. H., Beasley, J. C., & Jr, O. E. R. (2016). Carcass Type Affects Local Scavenger Guilds More than Habitat Connectivity. PLOS ONE, 11(2), e0147798. https://doi.org/10.1371/journal.pone.0147798

Onen, O. P., & Bassey, B. J. (2017). Biodiversity of City Dumpsites: What Future for the Environment? IOSR Journal Of Humanities And Social Science, 22(2), Article 2. https://doi.org/10.9790/0837-220201113119

Oro, D., Genovart, M., Tavecchia, G., Fowler, M. S., & Martínez-Abraín, A. (2013). Ecological and evolutionary implications of food subsidies from humans. Ecology Letters, 16(12), 1501–1514. https://doi.org/10.1111/ele.12187

Paswan, B. (2016). NATIONAL FAMILY HEALTH SURVEY (NFHS-4). International Institute for Population Sciences, Mumbai. Ministry of Health and Family Welfare, Government of India. http://rchiips.org/nfhs/nfhs-4Reports/India.pdf

Pattison, M. (Ed.). (2008). Poultry diseases (6th ed). Elsevier/Butterworth-Heinemann.

Pellow, D. N. (2004). Garbage Wars: The Struggle for Environmental Justice in Chicago. MIT Press.

Pickett, S. T. A., Cadenasso, M. L., Childers, D. L., McDonnell, M. J., & Zhou, W. (2016). Evolution and future of urban ecological science: Ecology in, of, and for the city. Ecosystem Health and Sustainability, 2(7), e01229. https://doi.org/10.1002/ehs2.1229

Pinault, D. (2008). Notes from the Fortune-Telling Parrot; Islam and the Struggle for Religious Pluralism in Pakistan; David Pinault. Equinox Publishing. https://www.equinoxpub.com/home/notes-fortune-telling-parrot/

Plaza, P. I., & Lambertucci, S. A. (2017). How are garbage dumps impacting vertebrate demography, health, and conservation? Global Ecology and Conservation, 12, 9–20. https://doi.org/10.1016/j.gecco.2017.08.002

Poore, J., & Nemecek, T. (2018). Reducing food’s environmental impacts through producers and consumers. Science, 360(6392), 987–992. https://doi.org/10.1126/science.aaq0216

Prakash, V., Pain, D. J., Cunningham, A. A., Donald, P. F., Prakash, N., Verma, A., Gargi, R., Sivakumar, S., & Rahmani, A. R. (2003). Catastrophic collapse of Indian white-backed Gyps bengalensis and long-billed Gyps indicus vulture populations. Biological Conservation, 109(3), 381–390. https://doi.org/10.1016/S0006-3207(02)00164-7

Prange, S., Gehrt, S. D., & Wiggers, E. P. (2003). Demographic Factors Contributing to High Raccoon Densities in Urban Landscapes. The Journal of Wildlife Management, 67(2), 324–333. https://doi.org/10.2307/3802774

Reese, J. (2005). Dogs and Dog Control in Developing Countries. State of the Animals 2005. https://www.wellbeingintlstudiesrepository.org/sota_2005/6

Reporter, S. (2021, December 21). 5,000 people attacked by dogs every day in Delhi: AAP. The Hindu. https://www.thehindu.com/news/cities/Delhi/5000-people-attacked-by-dogs-every-day-in-delhi-aap/article38007921.ece

Roser, M. (2013). Economic Growth. Our World in Data. https://ourworldindata.org/economic-growth

Rota, A. (2023). Gender and livestock: Tools for design. From: Livestock Thematic Papers Tools for project design. International Fund for Agricultural Development. https://www.ifad.org/documents/38714170/39148759/Gender+and+livestock.pdf/67c6dca9-4a11-4f53-931e-2ccb46105a3c

Samanta, I., Joardar, S. N., & Das, P. K. (2018). Biosecurity Strategies for Backyard Poultry: A Controlled Way for Safe Food Production. Food Control and Biosecurity, 481–517. https://doi.org/10.1016/B978-0-12-811445-2.00014-3

Satterthwaite, D., McGranahan, G., & Tacoli, C. (2010). Urbanization and its implications for food and farming. Philosophical Transactions of the Royal Society B: Biological Sciences, 365(1554), 2809–2820. https://doi.org/10.1098/rstb.2010.0136

Schell, C. J., Dyson, K., Fuentes, T. L., Des Roches, S., Harris, N. C., Miller, D. S., Woelfle-Erskine, C. A., & Lambert, M. R. (2020). The ecological and evolutionary consequences of systemic racism in urban environments. Science (New York, N.Y.), 369(6510), eaay4497. https://doi.org/10.1126/science.aay4497

Schwarzenbach, R. P., Egli, T., Hofstetter, T. B., von Gunten, U., & Wehrli, B. (2010). Global Water Pollution and Human Health. Annual Review of Environment and Resources, 35(1), 109–136. https://doi.org/10.1146/annurev-environ-100809-125342

Scoones, I. (2016). The Politics of Sustainability and Development. Annual Review of Environment and Resources, 41(1), 293–319. https://doi.org/10.1146/annurev-environ-110615-090039

Seress, G., & Liker, A. (2015). Habitat urbanization and its effects on birds. Acta Zoologica Academiae Scientiarum Hungaricae, 61(4), 373–408. https://doi.org/10.17109/AZH.61.4.373.2015

Shahmoradi, B. (2013). Collection of municipal solid waste in developing countries. International Journal of Environmental Studies, 70(6), 1013–1014. https://doi.org/10.1080/00207233.2013.853407

Sharma, K. (2022, December). MCD Elections 2022: Promises galore. ORF. https://www.orfonline.org/expert-speak/mcd-elections-2022-promises-galore/

Sharma, P., Maherchandani, S., Shringi, B. N., Kashyap, S. K., & Sundar, K. S. G. (2018). Temporal variations in patterns of Escherichia coli strain diversity and antimicrobial resistance in the migrant Egyptian vulture. Infection Ecology & Epidemiology, 8(1), 1450590. https://doi.org/10.1080/20008686.2018.1450590

Singh, M., Mollier, R. T., Paton, R. N., Pongener, N., Yadav, R., Singh, V., Katiyar, R., Kumar, R., Sonia, C., Bhatt, M., Babu, S., Rajkhowa, D. J., & Mishra, V. K. (2022). Backyard poultry farming with improved germplasm: Sustainable food production and nutritional security in fragile ecosystem. Frontiers in Sustainable Food Systems, 6. https://www.frontiersin.org/articles/10.3389/fsufs.2022.962268

Singh, S. (2021, August 20). The black kites of Ghazipur: In pandemic world, scavenging birds may foretell new maladies. Newslaundry. https://www.newslaundry.com/2021/08/20/the-black-kites-of-ghazipur-in-pandemic-world-scavenging-birds-may-foretell-new-maladies

Sinha, S. (1986). Economics vs Stigma: Socio-Economic Dynamics of Rural Leatherwork in UP. Economic and Political Weekly, 1061–1067.

Soga, M., & Gaston, K. J. (2016). Extinction of experience: The loss of human–nature interactions. Frontiers in Ecology and the Environment, 14(2), 94–101. https://doi.org/10.1002/fee.1225

Sorathiya, L. M., Fulsoundar, A. B., Tyagi, K. K., Patel, M. D., & Singh, R. R. (2014). Eco-friendly and modern methods of livestock waste recycling for enhancing farm profitability. International Journal of Recycling of Organic Waste in Agriculture, 3(1), 50. https://doi.org/10.1007/s40093-014-0050-6

Statistica. (2023). Topic: Waste generation worldwide. Statista. Retrieved November 8, 2022, from https://www.statista.com/topics/4983/waste-generation-worldwide/

Sumasgutner, P., Buij, R., McClure, C. J. W., Shaw, P., Dykstra, C. R., Kumar, N., & Rutz, C. (2021). Raptor research during the COVID-19 pandemic provides invaluable opportunities for conservation biology. Biological Conservation, 260, 109149. https://doi.org/10.1016/j.biocon.2021.109149

Talyan, V., Dahiya, R. P., & Sreekrishnan, T. R. (2008). State of municipal solid waste management in Delhi, the capital of India. Waste Management (New York, N.Y.), 28(7), 1276–1287. https://doi.org/10.1016/j.wasman.2007.05.017

Taneja, A. V. (2015). Saintly Animals: The Shifting Moral and Ecological Landscapes of North India. Comparative Studies of South Asia, Africa and the Middle East, 35(2), 204–221. https://doi.org/10.1215/1089201x-3138988

Thieme, O. (2013). Poultry Development Review. https://www.fao.org/3/i3531e/i3531e.pdf

Tscharntke, T., Clough, Y., Wanger, T. C., Jackson, L., Motzke, I., Perfecto, I., Vandermeer, J., & Whitbread, A. (2012). Global food security, biodiversity conservation and the future of agricultural intensification. Biological Conservation, 151(1), 53–59. https://doi.org/10.1016/j.biocon.2012.01.068

UNO. (2018). World Urbanization Prospects—Population Division—United Nations. https://population.un.org/wup/Publications/

Vaarst, M., Steenfeldt, S., & Horsted, K. (2015). Sustainable development perspectives of poultry production. World’s Poultry Science Journal, 71(4), 609–620. https://doi.org/10.1017/S0043933915002433

Vergara, S. E., & Tchobanoglous, G. (2012). Municipal Solid Waste and the Environment: A Global Perspective. Annual Review of Environment and Resources, 37(1), 277–309. https://doi.org/10.1146/annurev-environ-050511-122532

Vuorisalo, T., Talvitie, K., Kauhala, K., Bläuer, A., & Lahtinen, R. (2014). Urban red foxes (Vulpes vulpes L.) in Finland: A historical perspective. Landscape and Urban Planning, 124, 109–117. https://doi.org/10.1016/j.landurbplan.2013.12.002

Wang, X., Lu, X., Li, F., & Yang, G. (2014). Effects of Temperature and Carbon-Nitrogen (C/N) Ratio on the Performance of Anaerobic Co-Digestion of Dairy Manure, Chicken Manure and Rice Straw: Focusing on Ammonia Inhibition. PLOS ONE, 9(5), e97265. https://doi.org/10.1371/journal.pone.0097265

Wheat, C. H., Fitzpatrick, J. L., Rogell, B., & Temrin, H. (2019). Behavioural correlations of the domestication syndrome are decoupled in modern dog breeds. Nature Communications, 10(1), Article 1.

Wong, B. B. M., & Candolin, U. (2015). Behavioral responses to changing environments. Behavioral Ecology, 26(3), 665–673. https://doi.org/10.1093/beheco/aru183

Wong, J. T., de Bruyn, J., Bagnol, B., Grieve, H., Li, M., Pym, R., & Alders, R. G. (2017). Small-scale poultry and food security in resource-poor settings: A review. Global Food Security, 15, 43–52. https://doi.org/10.1016/j.gfs.2017.04.003

Woolhouse, M., Ward, M., van Bunnik, B., & Farrar, J. (2015). Antimicrobial resistance in humans, livestock and the wider environment. Philosophical Transactions of the Royal Society B: Biological Sciences, 370(1670), 20140083. https://doi.org/10.1098/rstb.2014.0083

Xu, Y., Gong, P., Wielstra, B., & Si, Y. (2016). Southward autumn migration of waterfowl facilitates cross-continental transmission of the highly pathogenic avian influenza H5N1 virus. Scientific Reports, 6(1), Article 1. https://doi.org/10.1038/srep30262

Yu, Y., & Zhang, W. (2016). Greenhouse gas emissions from solid waste in Beijing: The rising trend and the mitigation effects by management improvements. Waste Management & Research, 34(4), 368–377. https://doi.org/10.1177/0734242X16628982

